# B Cell Division Capacity in Germinal Centers Depends on Myc Transcript Stabilization Through m^6^A mRNA Methylation and IGF2BP3 Functions

**DOI:** 10.1101/2020.09.08.287433

**Authors:** Amalie C. Grenov, Lihee Moss, Sarit Edelheit, Ross Cordiner, Dominik Schmiedel, Adi Biram, Jacob H Hanna, Torben H Jensen, Schraga Schwartz, Ziv Shulman

## Abstract

Long-lasting immunity from pathogens depends on the generation of protective antibodies through the germinal center (GC) reaction. The *Myc* gene produces highly short-lived transcripts which are essential for generation of high-affinity antibodies. mRNA lifetime is regulated by N6-methyladenosine (m^6^A)-modification of mRNAs through METTL3 activity; however, the role of this machinery in the GC remains unclear. Here, we find that m^6^A-modification of mRNAs is required for GC maintenance through *Myc* mRNA stabilization by the atypical m^6^A-interactor, IGF2BP3. MYC expression, activation of MYC transcriptional programs and cell-cycle progression were diminished in METTL3-deficient GC B cells. METTL3 attenuated *Myc*-transcript decay and overexpression of MYC in METTL3-deficient GC B cells restored the GC reaction. IGF2BP3 which was induced by CD40-signaling, reinforced MYC expression and MYC-related gene programs in GC B cells. Our findings explain how GC responses are maintained through regulation of *Myc*-transcript lifetime and expose new targets for manipulation in MYC-driven lymphoma.

**One Sentence Summary:** Germinal centers depend on the m^6^A-machinery

## Introduction

The adaptive immune response has the capacity to generate long-lasting immunological memory that provides a rapid response to recurrent pathogen exposures (Ahmed and Gray, 1996; Cyster and Allen, 2019; Tarlinton and Good-Jacobson, 2013). The primary goal of vaccination is to induce the generation of long-lived antibody forming cells through the activation of the humoral arm of the adaptive immune response (Corcoran and Tarlinton, 2016; Ise and Kurosaki, 2019; Plotkin and Plotkin, 2008). High-affinity antibodies are secreted by plasma cells (PCs), which originate from germinal centers (GCs), microanatomical sites in lymphoid organs where B cells proliferate extensively and accumulate somatic hypermutations (SHM) in their immunoglobulin genes (Berek, 1992; Cyster, 2010; De Silva and Klein, 2015; Jacob et al., 1991; MacLennan, 1994; Shlomchik and Weisel, 2012; Victora and Nussenzweig, 2012). The selection of high-affinity clones for preferential expansion in the GC is mediated primarily by helper T cells (Crotty, 2011; Victora et al., 2010; Vinuesa and Cyster, 2011). In this process, the degree of T cell-derived signals induces corresponding expression levels of MYC in GC B cells which in turn activates downstream cell cycle- and metabolic gene programs (Calado et al., 2012; Chou et al., 2016; Dominguez-Sola et al., 2012; Ersching et al., 2017; Finkin et al., 2019; Gitlin et al., 2014, 2015; Monzón-Casanova et al., 2018). *Myc* is an immediate-early gene that rapidly produces mRNA transcripts with an extremely short half-life in response to mitogenic signals (Dani et al., 1984; Jones and Cole, 1987). How these short-lived transcripts support a continuous GC response is unclear. Ectopic MYC expression from a non-native locus (Rosa26) that lacks the natural regulatory elements, rescues GC formation in Myc-deficient B cells (Sander et al., 2012). These observations indicate that in addition to regulation of transcription activation, post-transcriptional events may play an important role in controlling MYC expression dynamics in the GC reaction.

Immune cell differentiation and acquisition of effector functions depend on extensive changes in gene expression that are regulated by transcriptional and post-transcriptional machineries (Turner and Díaz-Muñoz, 2018). The addition of a methyl group at position N6 on adenosines (m^6^A) is the most abundant post-transcriptional modification in mammalian mRNA (Shulman and Stern-Ginossar, 2020; Zaccara et al., 2019). This process is mediated by an mRNA modifying complex that includes METTL3, a methyltransferase that catalyzes the addition of m^6^A at consensus sites (Shulman and Stern-Ginossar, 2020; Zaccara et al., 2019). The functional consequence of the activity of this complex is determined by m^6^A-binding proteins that interact with the modified mRNA and modulate its half-life, location, splicing, and translation rate (Shi et al., 2017, 2019). Typically, m^6^A modifications promote mRNA destabilization, primarily through the binding of the m^6^A-interacting proteins, YTHDF1/2/3, in the cytosolic compartment (Du et al., 2016; Ivanova et al., 2017; Shi et al., 2017; Wang et al., 2014; Zaccara and Jaffrey, 2020). The functions of these m^6^A-binders in the adaptive immune response remain unknown. Conversely, IGF2BP1/2/3 (IMP1/2/3) is a family of mRNA binding proteins that stabilize mRNA transcripts and prolong their lifetime (Palanichamy et al., 2016; Ren et al., 2020). M^6^A modifications by METTL3 augment the binding capacity of IGF2BP paralogs to mRNAs resulting in the stabilization of several methylated transcripts, including DNA-replication- and cell-cycle-related genes (Huang et al., 2018). In particular, changes in transcript structure by m^6^A modifications and other supporting m^6^A-binding proteins may facilitate IGF2BP-mRNA interactions (Sun et al., 2019; Weidensdorfer et al., 2009; Youn et al., 2018; Zaccara et al., 2019). IGF2BP3 is the only paralog expressed in immune cells, yet its physiological functions within mammalian organisms remains unknown. Here, we examined how modulation of *Myc* mRNA lifetime through methylation controls B cell functions in GCs. We found that MYC-driven cell cycle progression in GC B cells depends on *Myc* mRNA stabilization through the activity of the m^6^A writer, METTL3, and the m^6^A interactor IGF2BP3. Thus, prolonging the lifetime of *Myc* mRNA in GC B cell-s through the m6A-machinery is essential for generation of effective and long-lasting humoral immune responses.

## Results

### Germinal center formation depends on METTL3

To examine whether METTL3 is required for generation and maintenance of the GC reaction, we crossed a Mettl3^fl/fl^ mouse strain to a mouse that expresses Cre recombinase under the B cell-specific CD19 promoter (abbreviated as CD19-Mettl3^fl/fl^); Mettl3^fl/+^ mice that express the Cre recombinase were used as controls. Since CD19 is expressed on all B cells during their generation in the bone marrow (BM), we first examined whether B cell development and establishment of the mature B cell compartment is defective in the CD19-Mettl3^fl/fl^ mouse model. Analysis of the different B cell populations in the BM and spleen did not reveal significant defects in B cell generation in CD19-Mettl3^fl/fl^ mice, and the pool of mature B cells in these mice was intact **(Fig. S1)**. Nonetheless, reduced levels of IgG1 and IgM antibodies were observed in the serum of the mutant strains under homeostatic conditions, suggesting a defect in the ability of Mettl3-deficient B cells to mount an effective immune response **(Fig. 1A)**. To examine whether METTL3 functions are required for the generation of GC B cells, control and CD19-Mettl3^fl/fl^ mice were immunized subcutaneously in the hind footpads with a hapten (4-hydroxy-3-nitrophenyl) coupled to keyhole limpet hemocyanin (NP-KLH) in alum, and the presence of GC B cells was examined in the popliteal lymph nodes (LNs) after 7 days. Flow cytometric analysis revealed that the frequency of GC B cells was 4-fold reduced in CD19-Mettl3^fl/fl^ mice compared to controls on day 7 of the response **(Fig. 1B)**. Furthermore, the frequencies of PCs and memory B cells in the LNs of the CD19-Mettl3^fl/fl^ mice were significantly lower compared to littermate controls **(Fig. 1B and S2A)**.

**Figure 1.**
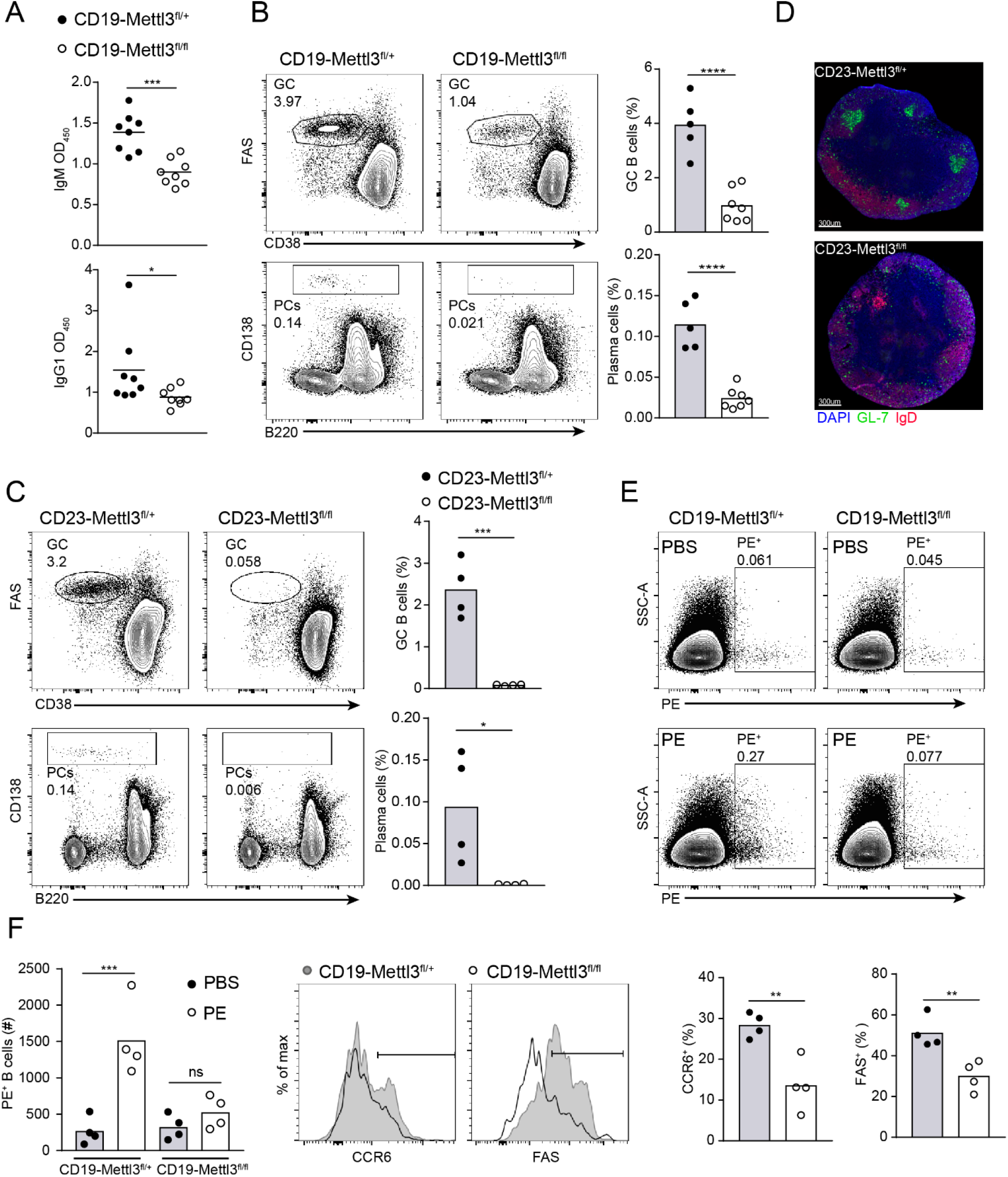
Mettl3-deficient B cells do not form germinal centers. (**A**) Serum immunoglobulin titers of IgM and IgG1 in CD19-Mettl3^fl/+^ or CD19-Mettl3^fl/fl^ of unmanipulated mice. 8-10 mice per condition, student’s paired *t* test between age-matched mice was used to determine significance (**B, C**) The frequency of GC B cells and PCs in control, CD19-Mettl3^fl/fl^ (B) or CD23-Mettl3^fl/fl^ (C) mice, 7 days after NP-KLH immunization. Each data point represents a single mouse and each column represents mean value. Pooled data from three experiments each with two or three mice per condition. (**D**) Representative immunostained slides of popliteal LNs derived from control or CD23-Mettl3^fl/fl^ mice. (**E and F**) Frequency and activation marker expression of PE-specific B cells in CD19-Mettl3^fl/+^ or CD19-Mettl3^fl/fl^ 3 days after immunization with PE. Each data point represents a single mouse and each column represents mean value. Pooled data from two experiments each with two mice per condition. \**P* < 0.05; \*\**P* < 0.01; \*\*\**P* <0.005, \*\*\*\**P* < 0.001; two-tailed Student’s *t* test (A-E) or one-way ANOVA corrected for multiple comparisons (Sidak) (F). NS, not significant.

To completely exclude the possibility that defects in GC formation are a result of aberrant B cell development in the BM of CD19-Mettl3^fl/fl^ animals (Zheng et al., 2020), we took an additional approach. CD23 is expressed on B cells in the spleen after B cell maturation and departure from the BM. Analysis of the B cell immune response in Mettl3^fl/fl^ mice that were crossed to a B cellspecific CD23-Cre mouse strain (CD23-Mettl3^fl/fl^) (Kwon et al., 2008) showed complete elimination of GC and PC formation 7 days after immunization **(Fig. 1C)**. Moreover, immunofluorescence analysis revealed that GC structures in CD23-Mettl3^fl/fl^ mice were absent **(Fig. 1D).** To determine whether early B cell activation or GC seeding were dependent on METTL3 functions we immunized CD19-Mettl3^fl/fl^ mice with Phycoerythrin (PE), a multi-epitope fluorescent antigen that allows detection of rare antigen-specific B cells in a polyclonal cell population at early time points (Pape et al., 2011). Flow cytometric analysis revealed a major defect in expansion of PE-specific B cells and in upregulation of the early activation markers, CCR6 and FAS in CD19-Mettl3^fl/fl^ mice 3 days after immunization **(Fig. 1E, F) (Schwickert et al., 2011).** These experiments demonstrate that METTL3 is essential for B cell functions prior to GC formation.

### METTL3 is required for sustained germinal center reaction

Activation-induced cytidine deaminase (AID, encoded by *Aicda*) is expressed in B cells after they encounter cognate antigen, but prior to GC seeding (Crouch et al., 2007; Roco et al., 2019; Rommel et al., 2013). To examine the functional role of METTL3 after the initial activation events, we crossed the Mettl3^fl/fl^ mice to an Aicda^Cre/+^ mouse strain (AID-Mettl3^fl/fl^). In these mice, serum IgG titers but not IgM titers were lower compared to controls **(Fig. S2B)**. As opposed to the observations in CD19-Mettl3^fl/fl^ mice, Mettl3 deletion at the time of AID expression did not affect GC seeding, and the frequencies of GC B cells, PCs and IgG1^+^ memory cells 7 days after immunization were intact **(Fig. 2A and S2C)**. However, after an additional week, the frequencies of these cell populations in AID-Mettl3^fl/fl^ mice were severely reduced compared to control mice **(Fig. 2A and S2C)**. The proportion of class-switched IgG1^+^ cells among Mettl3-deficient and control GC B cells was similar **(Fig. S2D)**. In some cases, deficiency in molecular machineries in B cell immune responses can only be detected under competitive pressure (Zaretsky et al., 2017). To examine whether METTL3 provides a competitive advantage in the GC reaction, we generated mixed BM chimeric mice that host control- and AID-Mettl3^fl/fl^-derived immune cells. In these mice, the frequency of Mettl3-deficient and proficient B cells in the GC was similar to that of naive B cells on day 7 after immunization **(Fig. 2B)**, suggesting that even under competitive pressure, METTL3 expression did not endow a competitive advantage at this time point. However, after an additional 7 and 14 days, the frequency of GC Mettl3-deficient B cells was severely reduced as observed under non-competitive conditions **(Fig. 2B)**. Furthermore, very few Mettl3-deficient B cells were detected in chronic GCs within the Peyer’s patches (PPs) of the chimeric mice **(Fig. S2E)**. Since GCs in PPs are driven by multiple gut-derived antigens, we conclude that the observed defects are not specific for any single immunogen. Collectively, these findings demonstrate that METTL3 functions are required for the persistence of B cells in the GC reaction in a cell intrinsic manner and regardless of competitive pressure.

**Figure 2.**
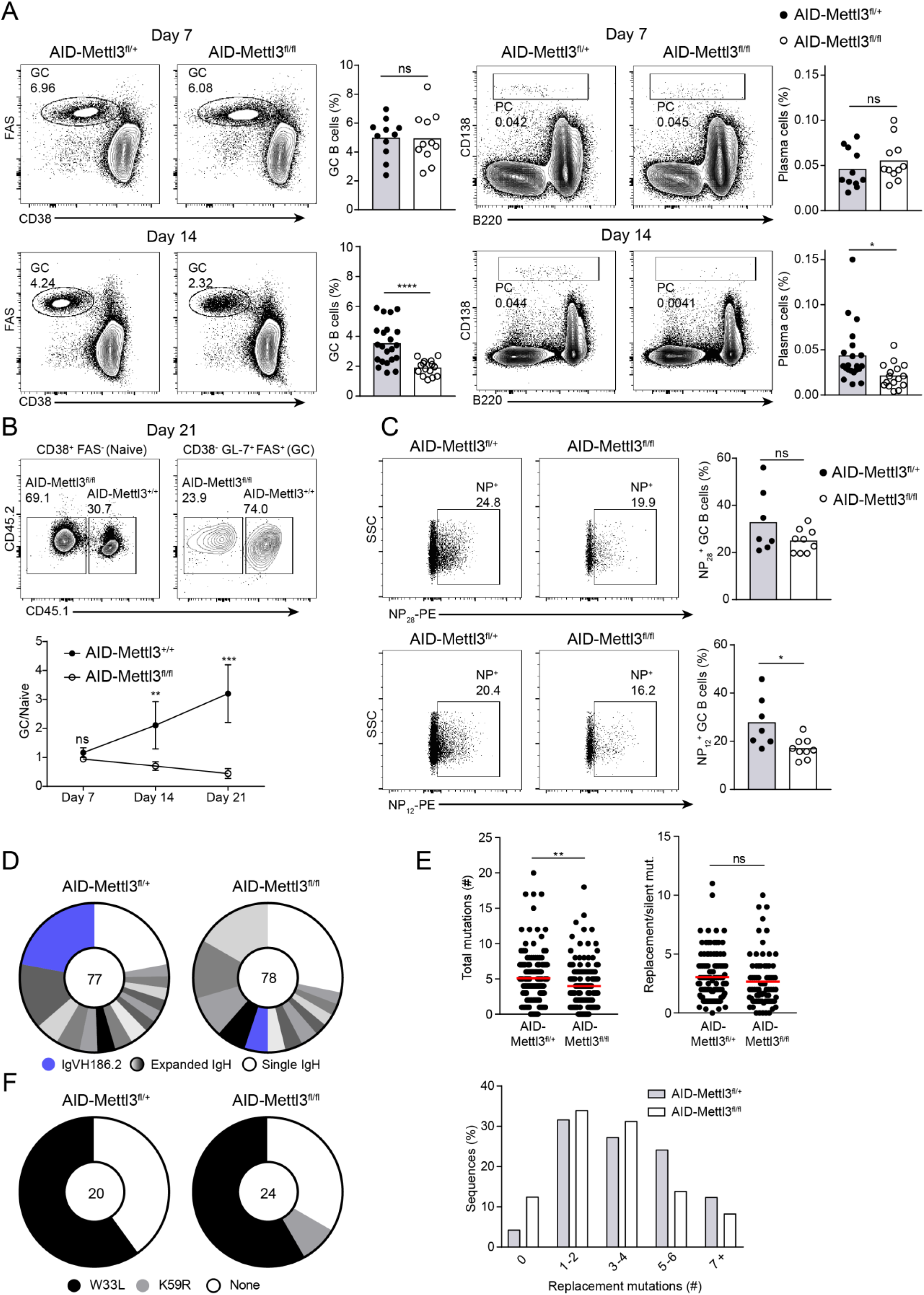
Durable germinal center reactions and proper clonal expansion requires METTL3. (**A**) The frequency of GC B cells and PCs in control and AID-Mettl3^fl/fl^ mice, 7 and 14 days after NP-KLH immunization. Data are pooled from 4-7 independent experiments with a total of 11-21 mice per group (**B**) Chimeric mice were generated by transfer of a mix of cells containing 70 % control BM cells (AID^cre/+^) and 30% AID-Mettl3^fl/fl^ BM cells. 1-3 weeks after immunization with NP-KLH, the proportion of control (CD45.1^+^) and Mettl3-deficient cells (CD45.2^+^) in the naïve and GC compartments in popliteal LNs were analyzed by flow cytometry. Data are pooled from 2-3 experiments with 2-3 mice per condition. Mean±SD are indicated. (**C**) The proportion of GC B cells that bind NP28-PE (total antigen-specific B cells) and cells that bind NP12-PE (high-affinity antigen-specific B cells), 21 days after immunization. Each data point represents a single mouse and columns represent mean values. Pooled data from three experiments, each with two or three mice per condition. (**D, E**) Single-cell analysis of immunoglobulin diversity (D), SHM frequency and amino acid replacement ratio (E), in IgG1^+^ GC B cells derived from control and AID-Mettl3^fl/fl^ mice, 21 days after immunization. Each slice in d represents a cluster of sequences with identical combinations of VDJs. Each data point in E represents a single immunoglobulin gene and lines represent mean values. (**F**) Frequencies of W33L and K59R mutations in VH186.2 of GC B cells as measured by bulk *Ighg* sequencing. Each slice represents a cluster of sequences with the indicated mutations. Two experiments each with one mouse per condition were performed and shown in a representative plot (D) or pooled data (E and F). \**P* < 0.05; \*\**P* < 0.01; \*\*\*\**P* < 0.001; two-tailed Student’s *t* test (A, C-D) or one-way ANOVA corrected for multiple comparisons (Sidak) (B). NS, not significant.

To determine whether the GCs in AID-Mettl3^fl/fl^ mice are functional and can support B cell SHM and selective clonal expansion, we first examined the presence of antigen-specific B cells in GCs by staining cells derived from immunized mice with PE-NP28 (which identifies total NP-specific B cells) or PE-NP12 (which identifies NP-specific B cells that express higher affinity BCRs). Flow cytometric analysis revealed that AID-Mettl3^fl/fl^ mice hosted slightly less NP-specific B cells in the GC; whereas, the proportion of B cells that bind NP12 was significantly reduced in these mice **(Fig. 2C)**. This result suggests that either AID-Mettl3^fl/fl^ B cells are defective in the acquisition of affinity-enhancing SHM in their immunoglobulin genes, or that subsequent clonal expansion is ineffective. To distinguish these possibilities, individual IgG1^+^ GC B cells were sorted from single LNs followed by *Igh* mRNA sequencing. This analysis revealed that the degree of immunoglobulin diversity within the GCs of control and AID-Mettl3^fl/fl^ mice was similar **(Fig. 2D)**. However, the typical clones that respond to NP in C57BL/6 mice (V_H_186.2) (Allen et al., 1987) were significantly less expanded in AID-Mettl3^fl/fl^ mice **(Fig. 2D)**. GC B cells in AID-Mettl3^fl/fl^ mice showed a small (1.3-fold) but significant reduction in the number of total mutations per cell **(Fig. 2E)**. Nonetheless, the ratio of replacement to silent mutations was similar; suggesting that positive selection of high affinity variants is intact in AID-Mettl3^fl/fl^ mice **(Fig. 2E)**. Accordingly, the few V_H_186.2 clonal members in the Mettl3-deficient GC B cells carried a mutation in the *Igh* V_H_186.2 (W33L or K59R), which serves as a marker for affinity-based selection (Allen et al., 1987; Furukawa et al., 1999). To more thoroughly probe whether acquisition of affinity-enhancing mutations is intact in the absence of METTL3, control and AID-Mettl3^fl/fl^ mice were immunized, and sorted Igλ^+^ IgG1^+^ GC B cells were subjected to *Igh* mRNA sequencing. Analysis of the V_H_186.2 mRNA transcripts revealed that Mettl3-deficient B cells show normal frequencies of replacement mutations that are associated with enhanced BCR affinity **(Fig. 2F)**. Thus, affinity-based clonal selection is intact in Mettl3-deficient GC B cells, whereas clonal expansion of the typically selected clones is suboptimal.

### Expression of dark zone-related gene programs depend on METTL3

Our findings demonstrate that METTL3 functions are required for maintenance of the GC size over time. The GC is composed of two compartments, the dark zone (DZ) where B cells acquire SHM and undergo clonal expansion, and the light zone (LZ) where affinity-based selection takes place (Allen et al., 2004, 2007; Victora and Nussenzweig, 2012). GCs of AID-Mettl3^fl/fl^ mice showed a higher proportion of LZ B cells (CXCR4^low^ CD86^high^) compared to control mice at day 14 of the response but not at day 7 **(Fig. 3A)**. To determine which molecular mechanisms depend on METTL3 functions in specific GC sites, LZ and DZ B cells were sorted from control and AID-Mettl3^fl/fl^ mice and subjected to RNA sequencing 7 and 14 days after immunization. Analysis of differential gene expression revealed that METTL3 plays a significant role in regulating gene expression in the DZ whereas, very small transcriptional differences were observed in the LZ at both time points **(Fig. 3B and S3A)**. In Mettl3-deficient DZ B cells, 226 genes showed increased expression, whereas 100 genes showed decreased expression compared to control cells (Fold change ≥ 1.5, pAdj = 0.1). Furthermore, whereas the transcriptome of control GC B cells showed expected differential gene expression between LZ and DZ, Mettl3-deficient GC B cells showed very small transcriptional differences between these compartments. **(Fig. 3C)**. Consistently, gene set enrichment analysis (GSEA) (Subramanian et al., 2005) revealed that Mettl3-deficient DZ B cells (sorted as CXCR4^high^ and CD86^low^) did not express typical DZ genes but rather showed increased expression of the typical LZ transcriptional program **(Fig. 3D)**. Transcription of genes that regulate critical biological processes in the DZ, including cell-cycle, DNA repair, and RNA metabolic pathways were significantly reduced in Mettl3-deficient DZ B cells **(Fig. 3E)**. In addition, GSEA revealed that cell cycle, MYC targets, and DNA repair-related genes were downregulated, whereas genes that are part of the apoptosis pathway were upregulated compared to control mice **(Fig. 3F, G)**. Together, these results indicate that although discernible DZ and LZ compartments that express zone-specific markers are formed independently of METTL3, this enzyme is required for proper expression of typical DZ-associated gene programs, including genes that are activated by MYC and cell cycle-related genes.

**Figure 3.**
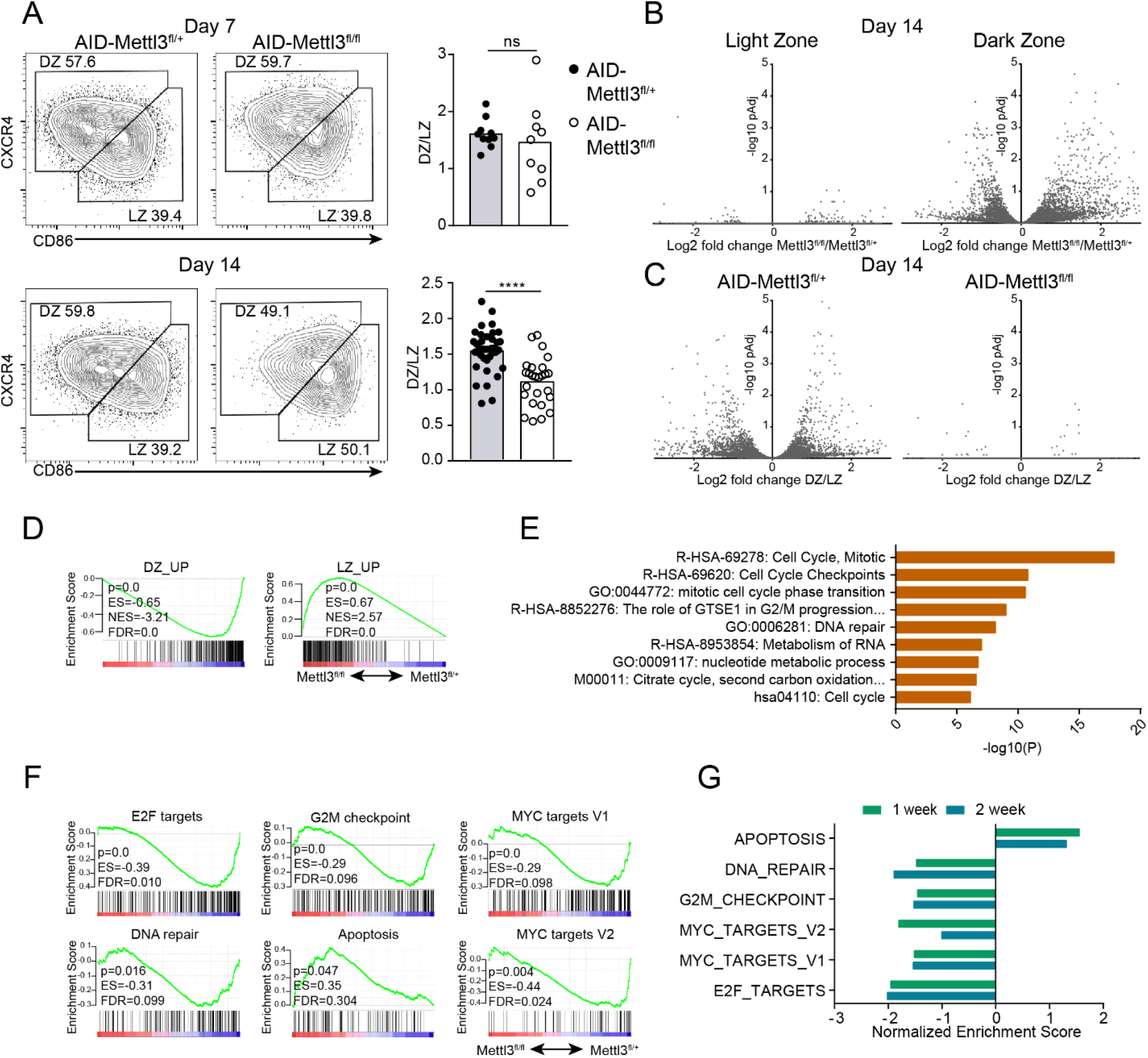
METTL3 is required for modulation of cell cycle-related genes in the dark zone. (**A**) The distribution of GC B cells to dark and light zone subsets in control and AID-Mettl3^fl/fl^ mice 7 and 14 days after NP-KLH immunization. Data pooled from 4-7 experiments with a total of 9-21 mice per group. (**B**) Differential gene expression in LZ and DZ GC B cells derived from mice in a (3 mice per condition), 14 days after immunization. (**C**) Differential gene expression between LZ and DZ GC B cells in either control or AID-Mettl3^fl/fl^ mice. (**D**) GSEA of DZ- and LZ-related genes in Mettl3-deficient GC B cells 14 days after immunization. p=P-value, ES=Enrichment Score, NES=Normalized Enrichment Score, FDR=False Discovery Rate. (**E**) Gene annotation analysis was performed on downregulated genes in Mettl3-deficient DZ B cells (fold change≥1.5, pAdj=0.1) 14 days after immunization, using Metascape online tool. (F) GSEA related to cell cycle progression, DNA-repair and apoptosis in DZ GC B cells 7 days after immunization. (G) Normalized gene enrichment scores for gene sets shown in f, 7 and 14 days after immunization. Two-tailed unpaired Student’s *t* test. \*\*\*\**P* < 0.001. NS, not significant.

### Efficient cell cycle progression of germinal center B cells depends on METTL3

To verify our transcriptomic findings in functional assays, we sought to measure cell division and cell cycle stages in GC B cells. To this end, we pulsed immunized mice with EdU, by intravenous injection, and examined the frequency of EdU^+^ cells after 2.5 hours. Flow cytometric analysis revealed that Mettl3-deficient GC B cells proliferated to a lesser degree (~5% fewer cells per 2.5 hours) than control mice during this short experimental period **(Fig. 4A)**. Co-labeling of DNA-incorporated EdU and total DNA revealed that significantly fewer cells were detected in the S phase of the cell cycle in Mettl3-deficient GC B cells **(Fig. 4B)**. These small differences in cell-cycle progression reflect a defect in proliferation capacity that can greatly affect the number of GC B cells over time.

**Figure 4.**
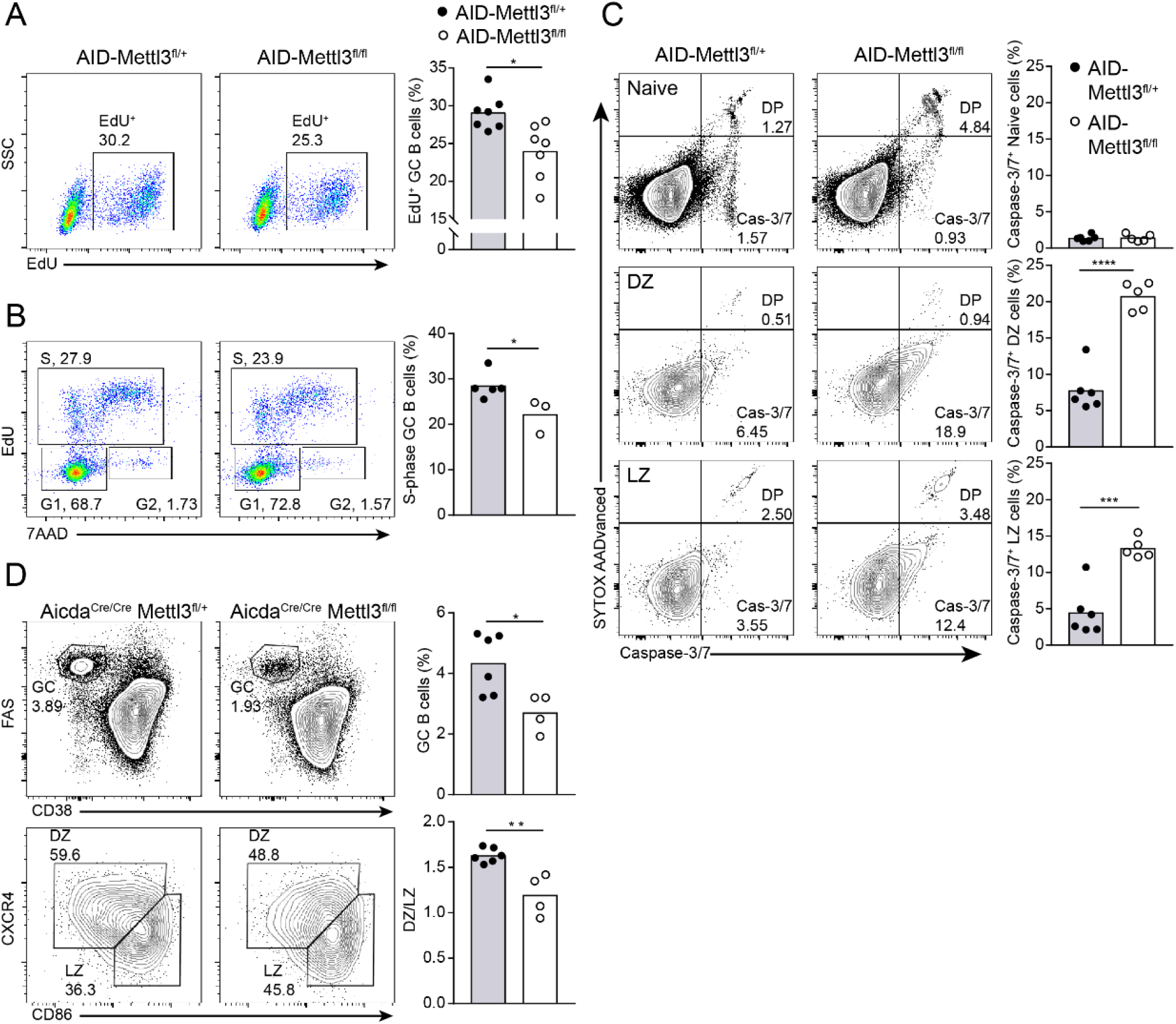
Effective cell cycle progression of germinal center B cells depends on METTL3. (**A, B**) Analysis of the different cell cycle stages by EdU incorporation and 7AAD DNA staining 14 days after NP-KLH immunization. Data are pooled from 2-3 experiments with a total of 3 to 7 mice per condition. **(C)** The frequency of apoptotic cells was measured by a caspase-3/7 activity assay and DNA content in naive, DZ and LZ B cells. (**D**) GC formation in *Aicda^Cre/Cre^* Mettl3^fl/+^ and *Aicda^Cre/Cre^* Mettl3^fl/fl^ mice 14 days after immunization. In c and d data are pooled from 2 experiments each with 2-3 mice per condition. Each data point represents a single mouse, and column heights represent mean values. Two-tailed Student’s *t* test. \**P* < 0.05;\*\**P* < 0.01; \*\*\**P* < 0.005; \*\*\*\**P* < 0.001. NS, not significant.

The transcriptomic analysis revealed that genes involved in apoptosis are upregulated in Mettl3-deficient GC B cells. To determine if the reduction in GC B cell frequencies is also a result of an increase in apoptotic events, in addition to reduced proliferation, we examined the activity of caspase-3/7. To distinguish between apoptotic and dead cells we co-labeled B cells with a DNA dye (SYTOX AADvanced, SAAD) and a cleavable peptide that labels cells with active caspase-3/7. SAAD and active caspase-3/7 positive signals label dead cells whereas SAAD negative and active caspase-3/7 markers label apoptotic cells. As expected, very few apoptotic cells were observed in the naive B cell compartment **(Fig. 4C)**. In contrast, apoptotic cells were clearly detected in the GC cell population and their frequency in Mettl3-deficient GC B cells was 3-fold higher than in the control in both the dark and light zones **(Fig. 4C)**. These results demonstrate that METTL3 activity prevents excess B cell apoptosis in the GC reaction.

AID enzymatic activity inserts mutations into the *Igh* genes that may result in the inability of GC B cells to express a functional B cell antigen receptor (BCR) (Mayer et al., 2017; Stewart et al., 2018). In addition, this enzyme acts as a major driver of off-target DNA damage events and chromosomal translocations in GC B cells (Casellas et al., 2016). Furthermore, AID activity induces DNA double strand breaks and METTL3 may play a role in the subsequent DNA repair mechanisms (Daniel and Nussenzweig, 2013; Zhang et al., 2020). To examine whether enhanced apoptosis and reduction in the frequency of GC B cells is a result of AID-mediated DNA damage or of defects in the subsequent DNA repair mechanism, we generated Aicda^cre/cre^ Mettl3^fl/+^ and Aicda^cre/cre^ Mettl3^fl/fl^ mice that lack expression of the AID enzyme. The deletion of AID did not increase the frequency of GC B cells in Aicda^cre/cre^ Mettl3^fl/fl^ immunized mice to resemble that of the control mice on day 14 after immunization, and the DZ to LZ ratio defect remained **(Fig. 4D).** These results suggest that deficiency in AID functions as well as in DNA repair of AID-mediated DNA damage cannot account for the decreased GC cell number in Mettl3-deficient mice. Collectively, these results suggest that METTL3 supports B cell persistence and survival in the GC primarily through regulation of proliferation mechanisms.

### *Myc* mRNA abundance and the positive B cell selection gene program are regulated by METTL3

The GC reaction and selection of high-affinity clones by T cells depend on MYC functions and activation of downstream gene transcription, promoting clonal expansion (Calado et al., 2012; Dominguez-Sola et al., 2012). The degree of B cell proliferation in the DZ is proportional to the initial magnitude of T cell help and level of MYC expression in the LZ (Finkin et al., 2019; Gitlin et al., 2014). GSEA revealed that GC-related gene programs were expressed at lower levels in Mettl3-deficient GC B cells compared to control mice two weeks after immunization **(Fig. 5A)**. To specifically characterize genes that are associated with positive B cell selection in the GC, we generated a “selection signature” based on a comparison between selected and non-selected GC B cells (Ersching et al., 2017). GSEA revealed a major reduction in the “selection signature” program in Mettl3-deficient GC B cells compared to control cells **(Fig. 5B)**. Consistent with these results, MYC downstream targets were significantly downregulated in GC B cells of Mettl3-deficient mice **(Fig. 3F)**. These findings are in agreement with the requirement for *Myc* transcription and expression of downstream genes for GC maintenance and proper selection of B cells for clonal expansion in response to T cell help and BCR signals (Calado et al., 2012; Dominguez-Sola et al., 2012; Finkin et al., 2019; Luo et al., 2018). RT-qPCR analysis of sorted GC B cells revealed reduced levels of *Myc* mRNA in Mettl3-deficient LZ B cells, and lower expression of MYC protein was observed in GC B cells derived from AID-Mettl3^fl/fl^ mice that express MYC-GFP compared to AID-Mettl3^fl/+^ litter mates **(Fig. 5C, D) (Huang et al., 2008).** To examine whether the maintenance of sufficient *Myc* transcripts through Mettl3 activity plays a dominant role in GC B cells we generated mice that overexpress MYC in Mettl3-deficient B cells (Sander et al., 2012). AID-Mettl3^fl/fl^ mice were bred to a mouse strain that carries the human *Myc* gene in the Rosa26 locus and a stop codon in front of it flanked by *LoxP* sites (R26^Myc/+^) (Sander et al., 2012). Age-matched AID-Mettl3^fl/+^, AID-Mettl3^fl/fl^ and AID-Mettl3^fl/fl^ R26^Myc/+^ mice were immunized with NP-KLH and the GC size was examined by flow cytometry after 14 days. Whereas the frequency of GC cells was smaller in AID-Mettl3^fl/fl^ mice compared to control, the AID-Mettl3^fl/fl^ R26^Myc/+^ mice showed normal frequencies of GC cells **(Fig. 5E)**. Thus, METTL3 functions regulate *Myc* transcript dosage and expression of downstream genes that promote GC maintenance and positive B cell selection for clonal expansion.

**Figure 5.**
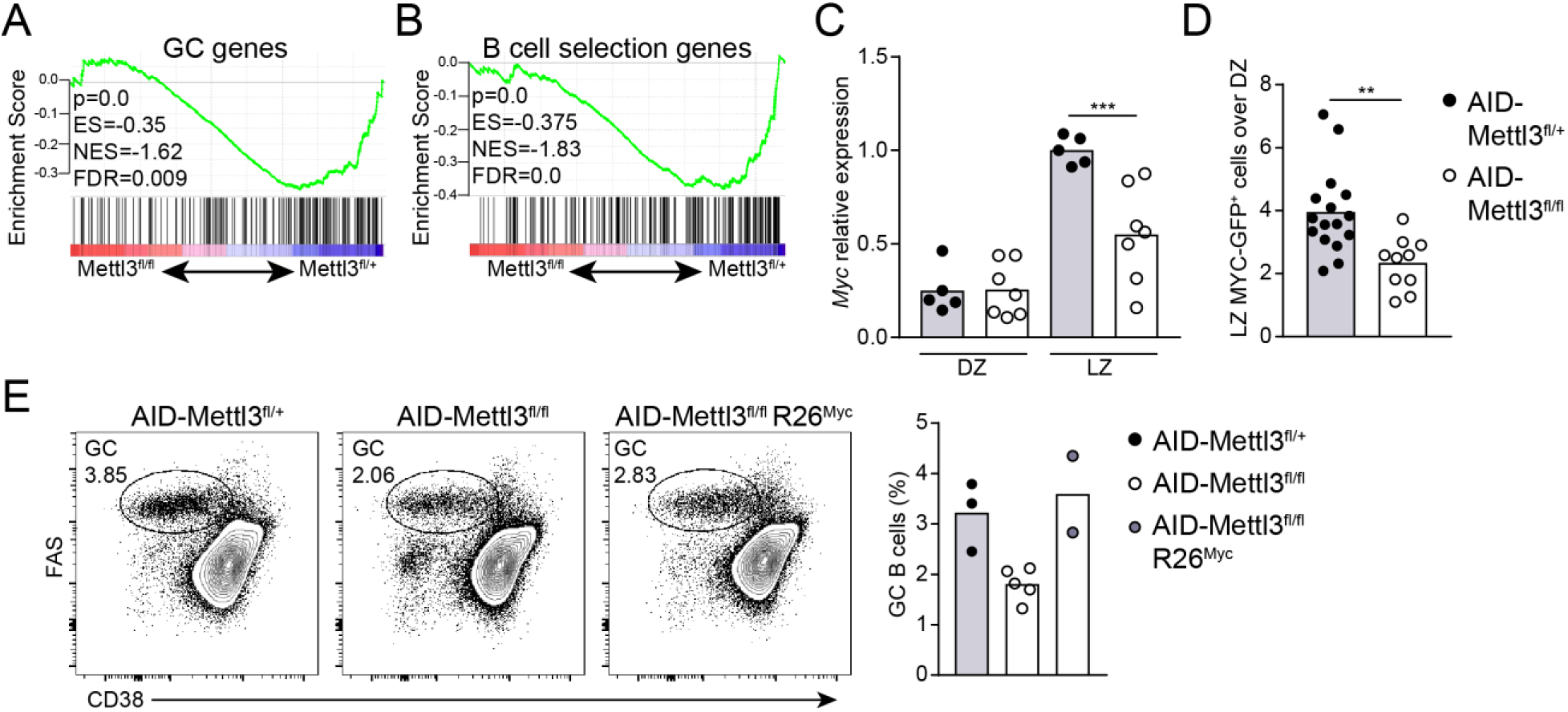
Effective MYC expression and the core germinal center genetic program depend on METTL3. (**A, B**) Enrichment of gene signatures related to GCs and B cell selection genes. (**C**) RT-qPCR analysis of relative amounts of *Myc* mRNA in GC B cells derived from AID-Mettl3^fl/+^ or AID-Mettl3^fl/fl^ 14 days after immunization. Pooled data from three experiments. Column height represents the mean. (**D**) MYC protein expression in GC B cells derived from cMYC-GFP AID-Mettl3^fl/+^ or cMYC-GFP AID-Mettl3^fl/fl^ immunized mice. Relative proportions of cMYC-GFP^+^ cells in GC LZ over DZ is shown. Pooled data from five experiments with a total of 10-16 mice per condition. (**E**) The frequency of GC cells was measured in age-matched AID-Mettl3^fl/+^, AID-Mettl3^fl/fl^ and AID-Mettl3^fl/fl^ R26^Myc/+^ 14 days after immunization. Pooled data from three experiments with a total of 2-5 mice per condition. One-way ANOVA corrected for multiple comparisons (Sidak)(C) or unpaired student’s *t* test (D). \*\**P* < 0.01; \*\*\**P* < 0.005. NS, not significant.

### METTL3 is required for *Myc* mRNA stabilization in germinal center B cells

The amount of mRNA in a given cell is determined by the rate of its transcription and degradation. To gain more insight into the abundance and persistence of *Myc* mRNA in the GC at the single cell level, we labeled *Myc* transcripts by single-molecule fluorescence *in situ* hybridization (smFISH) (Bahar Halpern and Itzkovitz, 2016). B cells in GC structures were identified by staining with Ki-67 probes and B220 antibody in lymph node slides derived from immunized mice **(Fig. S3B)**. Quantification of *Myc* transcript number per B cell revealed that in control mice 1.5-fold more single GC B cells expressed at least two mRNAs, compared to the corresponding Mettl3-deficient GC B cells **(Fig. 6A)**. The fewer *Myc* transcripts observed in total and single GC B cells can be a result of ineffective transcription or enhanced mRNA degradation rate. To distinguish between these two possibilities, we measured transcription rate through the visualization of *Myc* transcription start sites (TSs) in GC B cells by co-labeling *Myc* exons and introns in lymph node sections (Bahar Halpern and Itzkovitz, 2016; Halpern et al., 2015). The amount of nascent mRNAs within each TS site was determined based on the intensity of mature mRNA dots found outside the TS sites, (representing single mRNA molecules) and mRNA dot intensities in TSs (representing a few mRNA molecules that are being actively transcribed) (Bahar Halpern and Itzkovitz, 2016; Halpern et al., 2015). By integrative analysis of the nascent and mature mRNA molecules, we found that the rate of *Myc* transcription *in situ* in Mettl3-deficient GC B cells was similar to controls **(Fig. 6B, C)**. Nonetheless, the abundance of mature *Myc* transcripts was lower in Mettl3-deficient GC B cells that actively transcribed *Myc,* demonstrating that the degradation rate was higher in AID-Mettl3^fl/fl^ mice **(Fig. 6B, C)**. We conclude that the lower amounts of *Myc* mRNA observed by RT-qPCR and smFISH as well as the defect in MYC-downstream gene expression in Mettl3-deficient GC B cells, was not as a result of direct or indirect defects on *Myc* transcription but rather of enhanced *Myc* mRNA degradation rate.

**Figure 6.**
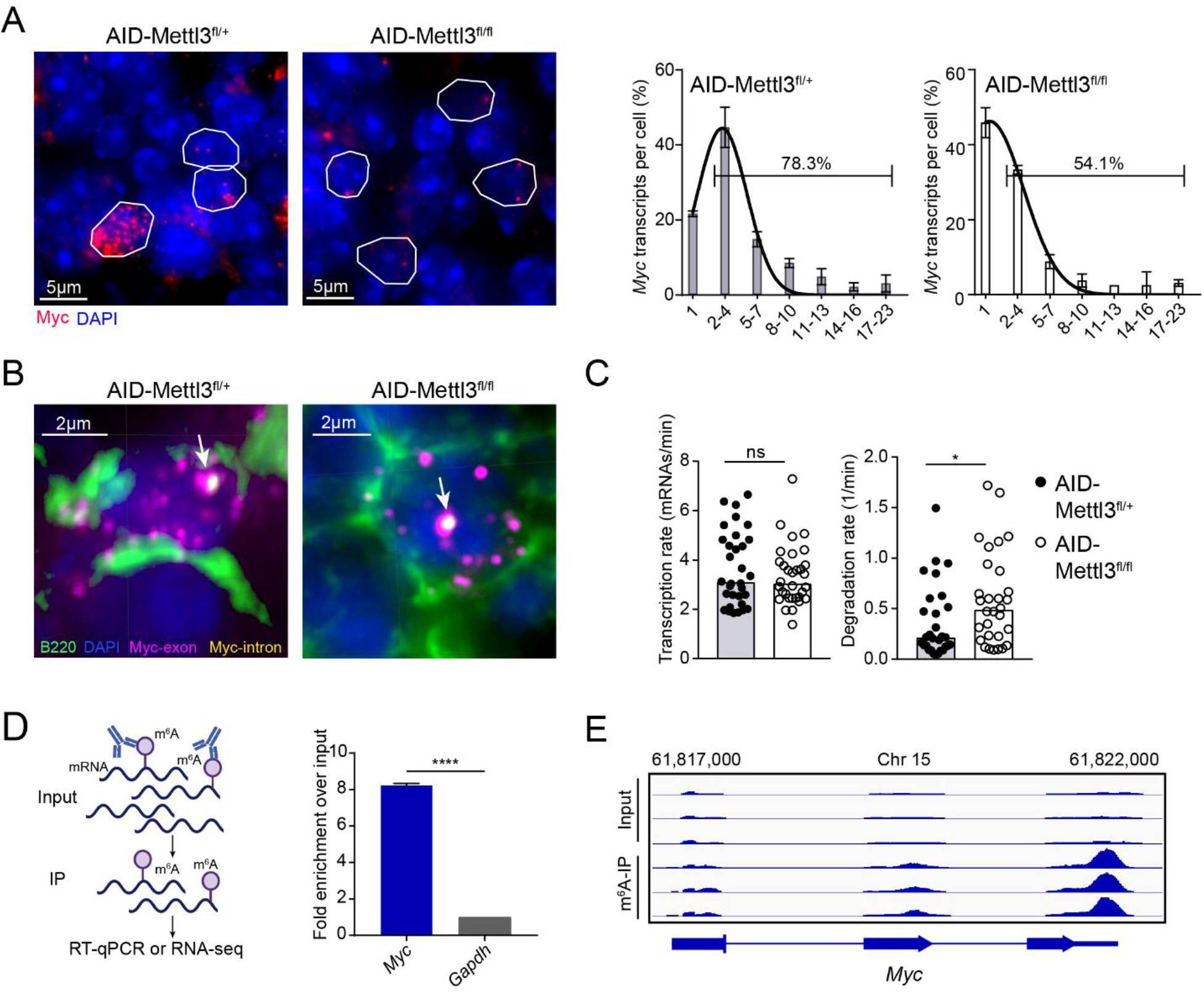
METTL3 prolongs the lifetime of methylated *Myc* transcripts in germinal center B cells. (**A-C**) Myc mRNA abundance (A), transcription and degradation rates (B, C) in GC B cells by smFISH. GCs were identified by B220 antibody staining (green) and Ki-67 (Fig. S4E) smFISH probes. Red (A) and Magenta (B) dots are *Myc* mRNA detected using a probe library targeting the exons. Yellow dots (B) are pre-mRNAs detected using a probe library targeting the introns. Active TSs are identified by dual colour labeling. Representative images from 2 mice, 3-5 GCs and 31-35 cells per condition are shown and *Myc* transcripts per cell (A), transcription and degradation rates (B) are summarized in the graphs. Column height represents mean+SD (A) or median+SD (C). (**D**) *Myc* and *Gapdh* mRNA methylation in LPS-stimulated B cells was measured by m^6^A-RIP followed by RT-qPCR. Fold enrichment of *Myc* mRNA in pulldown over input fraction normalized to unmethylated *Gapdh* mRNA is shown. Pooled data from two replicates. (**E**) *Myc* methylation sites analysis by pull down as in (D) followed by high-throughput sequencing. Mapped reads to *Myc* from the input and m6A-IP fraction are indicated. Unpaired student’s *t* test was used to determine statistical significance in C and D and Two-way ANOVA was used in A, with *P* of the interaction = 0.0001. \**P* < 0.05; \*\*\*\**P* < 0.001; NS, not significant.

METTL3 is a methyltransferase that catalyzes addition of m^6^A on mRNA transcripts at consensus sites. To examine whether *Myc* mRNA carries m^6^A modifications in B cells, we performed immunoprecipitation of methylated mRNA (m^6^A-IP) followed by RT-qPCR or mRNA sequencing **(Fig. 6D)**. Since we could not obtain sufficient amounts of mRNA from GC B cells for this analysis, we examined primary B cells that were stimulated with LPS *ex vivo.* As opposed to their severe defect in responding to stimulation *in vivo,* Mettl3-deficient B cells treated with LPS *ex vivo* proliferated only slightly less than control cells. In this setting, the dividing cells were found to express METTL3, suggesting that a few cells in which *Mettl3* was not deleted, responded to the LPS stimulation and proliferated extensively **(Fig. S4A, B)**. This observation is consistent with previous reports that show a strong selection pressure to maintain METTL3 expression *in vitro* (Cheng et al., 2019b; Liu et al., 2014; Schwartz et al., 2014). M^6^A-IP followed by RT-qPCR of WT B cells that were stimulated with LPS for 3 days revealed that *Myc* mRNA was highly enriched in the pulldown fraction compared to control unmethylated mRNA *(Gapdh*) **(Fig. 6D)**. To further validate the methylation of *Myc* transcripts we performed m^6^A-IP followed by high throughput sequencing. M^6^A-seq on mRNA derived from Mettl3-deficient B cells that were stimulated with LPS, yielded peak over input scores that were reduced to 60% of control **(Fig. S5)**. The m^6^A sites that were specifically detected in control cells were enriched around m^6^A consensus motifs, 3’UTRs and around stop codons **(Fig. S5)**. Notably, *Myc* mRNAs was found to carry methylations at sites that are close to the stop-codon region **(Fig. 6E).** Thus, *Myc* mRNA is methylated in B cells, however, *ex vivo*-activated B cells are not an ideal model for analysis of Mettl3 functions.

### IGF2BP3 but not YTHDF2 is required for effective activation of MYC-related gene programs

The m^6^A modification of mRNAs through METTL3 activity can be recognized by YTHDF mRNA binders that control transcript lifetime and translation rate (Zaccara et al., 2019). As previously shown in cell lines (Patil et al., 2018; Zaccara and Jaffrey, 2020), among the different YTHDF paralogs, *Ythdf2* mRNA was most highly expressed in GC B cells (**Fig. S3C)**. This specific family member is associated with transcript destabilization and degradation (Du et al., 2016; Wang et al., 2014; Zaccara and Jaffrey, 2020). To examine if METTL3 functions in GC B cells depend on YTHDF2, we produced CD23-Ythdf2^fl/fl^ and CD23-Ythdf2^fl/+^ mice and immunized them with NP-KLH (Ivanova et al., 2017). Flow cytometric analysis revealed that CD23-Ythdf2^fl/fl^ mice showed a smaller frequency of GC B cells on day 14 after immunization compared to control mice **(Fig. 7A)**. To examine whether MYC is involved in the function of YTHDF2, we sorted GC B cells and examined *Myc* expression and the global transcriptome by RT-qPCR and RNA sequencing, respectively. As opposed to the results obtained in Mettl3-deficient mice, the amount of *Myc* transcripts was higher in Ythdf2-deficient GC B cells than in control cells **(Fig. 7B)**. In addition, MYC-responsive genes, the core GC genes, and gene programs associated with selection of high affinity clones, were not altered **(Fig. 7C and S3D)**. These results suggest that YTHDF2 plays a role in the GC, however, these effects are not mediated by downmodulation of MYC- and the core GC-related genes. We conclude that the reduced expression of MYC and MYC-induced genes in Mettl3-deficient B cells is not a result of the inability of YTHDF2 to directly bind methylated *Myc* mRNAs. Furthermore, these findings demonstrate that not all perturbations in the m^6^A-machinery that affect the size of the GC necessarily affect MYC functions, and the core of the GC-related genes.

**Figure 7.**
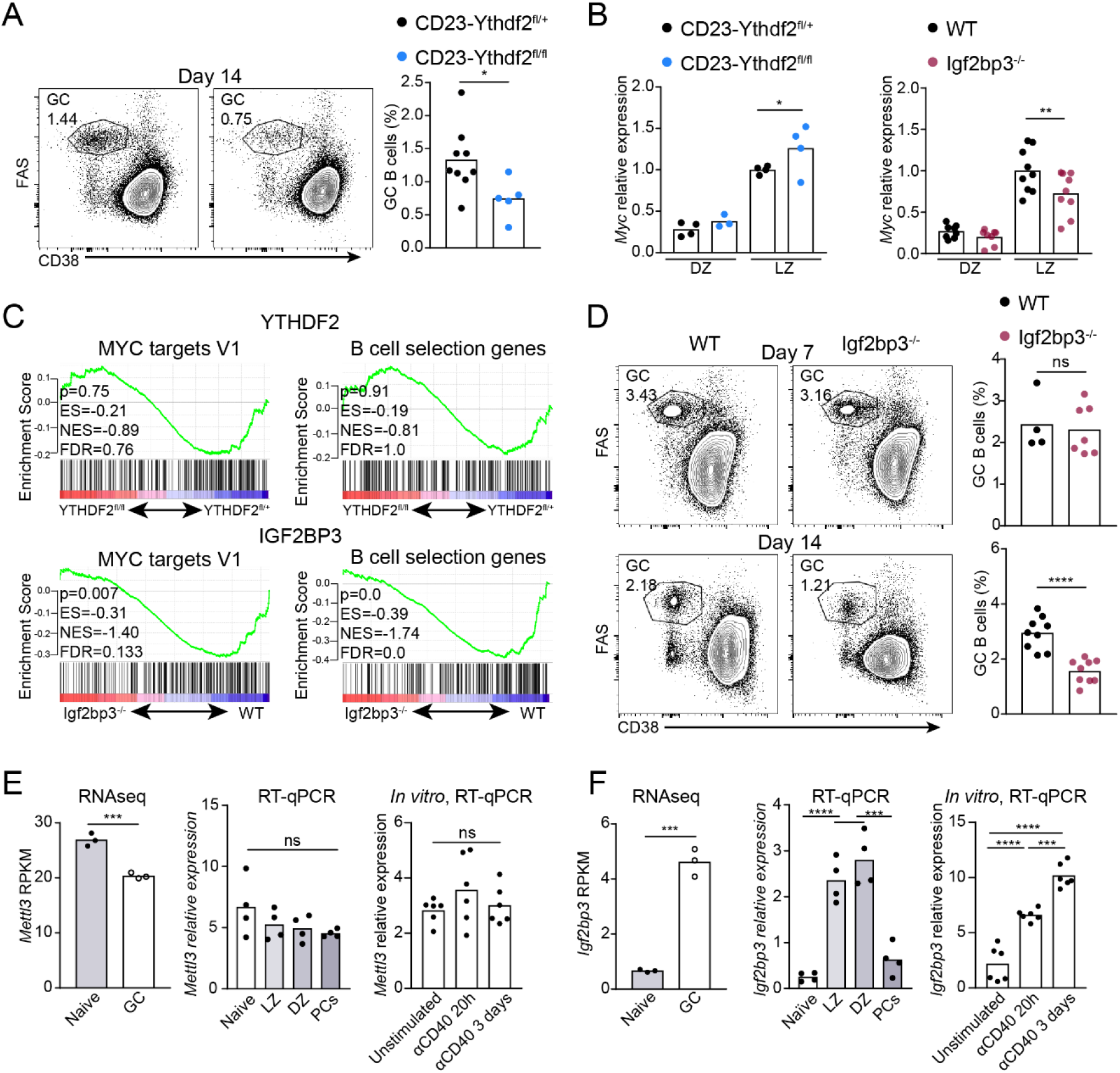
IGF2BP3 is required for effective MYC expression and activation of MYC targets. (**A**) Frequency of GC B cells and PCs in control, and AID-Ythdf2^fl/fl^ mice, 14 days after NP-KLH immunization. Data pooled from three experiments with 5-9 mice per condition. (**B**) Relative *Myc* mRNA expression in GC B cells derived from AID-Ythdf2^fl/+^, AID-Ythdf2^fl/fl^, WT or Igf2bp3^-/-^ mice 14 days after immunization. Data pooled from 2-3 experiments with a total of 3 to 9 mice per condition. (**C**) GSEA of MYC targets and GC B cell selection-related genes in AID-Ythdf2^fl/+^, AID-Ythdf2^fl/fl^, WT and Igf2bp3-deficient GC B cells. Gene expression differences based on 2-4 mice per condition. (**D**) The frequency of GC B cells and PCs in WT, and Igf2bp3^-/-^ mice, 7 and 14 days after NP-KLH immunization. Data pooled from two to three experiments each with 2-3 mice per condition. (**E, F**) Relative expression of *Mettl3* (E) and *Igf2bp3* (F) mRNAs as measured by RNA-seq data (left panels, GSE98086) and RTqPCR (middle and right panels). Data pooled from 3-9 mice (naive and GC cells) or two mice with three samples in each (CD40-stimulation). Expression by RT-qPCR was determined relative to housekeeping genes *Ubc* and *Ywhaz*. Two-tailed Student’s *t* test (A, D and E left panels) or one-way ANOVA corrected for multiple comparisons (Sidak) (B and E right panels). \**P* < 0.05; \*\**P* < 0.01; \*\*\**P* < 0.005; \*\*\*\**P* < 0.001. NS, not significant.

IGF2BPs are a family of mRNA binding proteins that increase the mRNA lifetime of target genes and whose binding capacities are enhanced by m^6^A methylation (Huang et al., 2018; Palanichamy et al., 2016; Ren et al., 2020; Samuels et al., 2020). IGF2BP3 binds the 3’-UTR of *Myc* mRNA in cell lines and is the only paralog that is expressed in immune cells (Palanichamy et al., 2016). Since m^6^A enhances IGF2BP3-mediated stabilization of *Myc* mRNA (Huang et al., 2018), we sought to examine the role of IGF2BP3 in GCs. The size of the GC compartment 7 days after immunization in Igf2bp3-deficient mice was similar to that of WT mice; however, the frequency of GC B cells decreased after an additional 7 days **(Fig. 7D)**. This defect was B cell-specific, as a similar phenotype was observed in chimeric mice hosting both WT and *Igf2bp3^-/-^* immune cells **(Fig. S3E)**. In addition, *Myc* mRNA levels were significantly reduced in sorted Igf2bp3-deficient LZ GC B cells **(Fig. 7B)**. To examine the effect of IGF2BP3 on MYC-responsive genes, GC B cells were isolated from mice 14 days after immunization and subjected to RNA sequencing followed by GSEA. Significant downregulation of MYC responsive genes, GC-related genes, and B cell selection signatures, as well as a loss of DZ transcriptional signature, were observed in Igf2bp3-deficient mice **(Fig. 7C and S3F)**. Collectively, we conclude that adequate expression of *Myc* mRNA and proper expression of MYC downstream targets in GC B cells depend on IGF2BP3.

Next, we examined whether the m^6^A-machinery is subjected to regulation during the B cell immune response. Analysis of RNA-seq data, as well as qRT-PCR revealed that METTL3 is expressed to a similar extent in the naïve and GC B cells, suggesting that regulation of m^6^A functions is not at the methyltransferase level **(Fig. 7E)**. Indeed, B cell activation through CD40 triggering did not induce changes in METTL3 mRNA levels **(Fig. 7E)**. In contrast, the amount of IGF2BP3 in GC B cells was ~10-fold higher compared to their naïve counterparts and CD40 triggering was sufficient to induce its expression in B cells **(Fig. 7F)**. These findings indicate that mRNA methylation by METTL3 is a constitutive process in B cells, whereas its downstream functions are regulated by extracellular signals that induce the expression of the m^6^A interactors such as IGF2BP3.

## Discussion

MYC is essential for generation and maintenance of the GC reaction and for preferential clonal expansion of high-affinity B cell clones; however, *Myc* transcripts have a very short half-life and are expressed in small amounts in the GC (Calado et al., 2012; Dani et al., 1984; Dominguez-Sola et al., 2012; Finkin et al., 2019; Jones and Cole, 1987). Here, we demonstrate that the m^6^A-machinery has a critical role in the stabilization of *Myc* transcripts during the GC reaction, wherein MYC expression in B cells is a limiting factor (Finkin et al., 2019). Altogether, we show that Myc lifetime and B cell fate are regulated by expression of IGF2BP3 through CD40 signalling, linking an extracellular signal, such as T cell help, to fate decision in B cells through the m^6^A machinery. In a broader view, our findings explain how short-lived transcripts that determine cell fate, such as *Myc,* are maintained for optimal gene expression and promote fate decisions under physiological conditions.

MYC expression is induced upon B cell interaction with antigen, but after 2 days, the amount of MYC protein is rapidly decreased followed by a limited re-expression in the fully formed GC (Calado et al., 2012; Dominguez-Sola et al., 2012). The MYC expression dynamics in B cells is consistent with our findings that demonstrate a requirement for METTL3 at the early and late stages of the response but not during initial AID expression at the time of pre-GC events (Calado et al., 2012; Dominguez-Sola et al., 2012). In particular, *Myc-inducing* factors such as cognate antigen and T cell help signals are limited at later stages of the response when METTL3 functions and IGF2BP3 are essential for maintaining the GC reaction. Although MYC expression is also associated with selection of high-affinity clones in the GC through competition for T cell help (Dominguez-Sola et al., 2012), the frequency of Mettl3-deficient B cells in GCs declined over time independently of competition with Mettl3-expressing cells, indicating that the m^6^A-machinery primarily plays a role in global B cell proliferation machineries in the GC rather than specifically in the positive clonal selection process. In support of this conclusion, we found that the acquisition of affinity-enhancing mutations in the GC is METTL3-independent whereas effective proliferation and proper expression of the GC core genes required METTL3 functions. Our findings are consistent with a model wherein T cells are able to discern between low and high affinity clones in the GC independently of m^6^A modifications in B cells, however, effective global and preferential clonal expansion in the GC dark zone depends on m^6^A mRNA modifications including *Myc* methylation. This strategy of GC cell maintenance is different from the HSC self-renewal process, for which METTL3 functions are not required for homeostatic MYC-independent proliferation (Cheng et al., 2019b; Lee et al., 2019).

The outcome of mRNA methylation depends on interactions with reader proteins. Typically, the deletion of *Mettl3* or *Ythdf2* in different cell types leads to an increase in the abundance of many target transcripts (Ivanova et al., 2017; Zaccara et al., 2019). In contrast, *in vitro* analyses revealed that MYC responsive genes are downregulated upon deletion of *Mettl3* or its binding partner *Mettl14* as a result of destabilization of *Myc* transcripts; and that these effects depend on methylation sites (Cheng et al., 2019a, 2019b; Huang et al., 2018; Vu et al., 2017; Weng et al., 2018; Zhao et al., 2020). Indeed, m^6^A pull down experiments revealed that *Myc* transcripts were highly methylated in activated B cells. Typically, transcript lifetime is measured in vitro by blocking transcription followed by quantification of mRNA abundance over time (Huang et al., 2018; Liu et al., 2014). Measurements of mRNA lifetime in cells that do not survive in culture without additional stimulation, such as GC B cells, is nearly impossible. To overcome this limitation we quantified single mRNA molecules *in situ* within the GC B cells (Bahar Halpern and Itzkovitz, 2016); we directly demonstrate that METTL3 in immune cells is required, specifically, for the stabilization of *Myc* under physiological conditions. Most importantly, our direct quantification of *Myc* mRNAs in GC B cells, demonstrates that the decline in transcript abundance is not a result of indirect effects such as reduction in expression of a transcription factor that controls *Myc* transcript dosage.

In non-mammalian model organisms and cell lines, IGF2BP paralogs enhance the stability of several mRNAs including *Myc* transcripts, and m^6^A has been shown to reinforce these mRNA-protein interactions (Huang et al., 2018; Palanichamy et al., 2016; Ren et al., 2020; Samuels et al., 2020). Our findings add a new role for this mRNA stabilization machinery in immune cells and demonstrate how IGF2BP3 controls cell fate in a mammalian organism. Whether IGF2BP3 functions depend only on its direct binding to m^6^A (Huang et al., 2018) or additional indirect events, such as m^6^A-dependent changes in mRNA structures or interactions with other typical m^6^A readers, such as YTHDF2, require additional investigation (Sun et al., 2019; Weidensdorfer et al., 2009; Youn et al., 2018; Zaccara et al., 2019). In addition, we revealed that YTHDF2 depletion in B cells reduces the GC size, however, and in agreement with previous reports (Huang et al., 2018; Wang et al., 2018), the abundance of *Myc* transcripts increased in Ythdf2-deficient B cells and there was no significant effect on the expression of MYC-downstream genes. These findings demonstrate that maintenance of sufficient *Myc* transcript levels through METTL3 functions does not depend on the dominant YTHDF family member in the GC. Whereas IGF2BP3 and YTHDF2 play critical roles within the established GC, which m^6^A-binders regulate early B cell activation *in vivo* remains to be determined.

CD40 signaling promotes the expression of both MYC (Luo et al., 2018) and IGF2BP3 in B cells suggesting that the interactions with T cells in the GC may induce the transcription of short lived *Myc* mRNA together with a protein that prolongs its lifetime. We did not find changes in METTL3 expression during the B cell immune response or after CD40 activation whereas the expression IGF2BP3 changed dramatically. These observations suggest that extracellular signals affect the outcome mRNA methylation through regulation of specific binder expression rather than METTL3-mediated methylation events per se.

Collectively, our findings define how the m^6^A-machinery supports a central process in the adaptive immune response through regulation of mRNA lifetime. Dysregulation of MYC expression drives the development of several cancers including the GC-derived Burkitt’s lymphoma (Sander et al., 2012). This study suggests that components of the m^6^A-machinery can serve as a target for manipulation of the adaptive arm of the immune system in pathological conditions such as in MYC-driven lymphoma and antibody-mediated autoimmune diseases.

## Materials and Methods

### Mice

Mettl3^fl/fl^ mice were generated and provided by Y. Hanna and crossed to Cre expressing mice and B cell specific Cd23cre mice were generated and provided by M. Busslinger (IMP, Vienna). Aicda^Cre/+^, Cd19-Cre, MYC-GFP and Igf2bp3^-/-^ mice were purchased from Jackson Laboratories. Ythdf2^fl/fl^ were provided by Donal O’Carroll (University of Edinburgh, UK). MYC-GFP expressing mice were bred to mice carrying Aicda^Cre^ and Mettl3^fl/fl^ alleles. Wild-type mice (C57BL/6) were purchased from Envigo. WT and transgenic mice were used at the age of 6-12 weeks. To avoid age variations in experiments that include a small number of mice that carry R26flox-stop-flox Myc gene, only age-matched mice (~8 weeks) were used. All experiments on mice were approved by the Weizmann Institute Animal Care and Use Committee.

### Chimeric mice

Chimeric mice were generated by irradiating host mice with 950rad followed by injection of fresh BM cells. For the generation of mixed chimeras, CD45.2^+^ hosts were reconstituted with a mixture of BM cells derived from Aicda^Cre/+^ CD45.1^+^ (50%) and Aicda^Cre/+^ Mettl3^fl/fl^ CD45.2^+^ (50%) mice or from CD45.1^+^ (50%) and Igf2bp3^-/-^ CD45.2^+^ (50%) mice. Experiments with chimeric mice were performed 8 weeks after BM transplantation.

### Immunizations

NP-KLH (BioSearch Technologies, CA, USA) was prepared with Alum and PBS at a final concentration of 0.4mg/ml. Mice received a single administration to each footpad of 25μl (10μg NP-KLH). For immunizations with PE, 10μg PE (Molecular Probes) in alum was injected.

### In vivo EdU proliferation and apoptosis measurements

Immunized mice were injected intravenously with 2 mg of the nucleoside analog EdU (Molecular Probes) in PBS. After 2.5 h, popliteal LNs were dissected and isolated LN cells were stained for surface antigens as described, followed by EdU detection using the Click-iT EdU Alexa Fluor 647 Flow Cytometry Assay Kit (Molecular Probes) according to manufacturer’s instructions. 7AAD was from BD Biosciences and added according to manufacturer’s instructions.

For measurements of apoptosis and cell death, popliteal LNs from immunized mice were isolated and single cell suspensions were stained for GC surface antigens and analyzed for apoptotic events using CellEvent™ Caspase-3/7 Green Flow Cytometry Assay Kit (Molecular Cell Probes) according to manufacturer’s instructions.

### Flow cytometry

Popliteal LNs were removed, washed in cold PBS and forced through a mesh into PBS containing 2% fetal calf serum and 1mM EDTA to create single cell suspensions. Cell suspensions from PPs were similarly prepared by excising PPs from small intestines pre-washed with ice-cold PBS to remove fecal content. To block Fc receptors, washed cells were incubated with 2μg/ml anti-16/32 (TruStain FcX, BioLegend) for 5-10min prior to antibody staining. Cells were subsequently incubated with fluorescently labeled antibodies (Table S1) for 30min on ice. Intracellular antibody staining for METTL3 was performed after fixation and permeabilization with Fixation/Permeabilization Solution Kit (BD biosciences).

GC cells were gated as live/single, B220^+^ CD38^-^ FAS^+^ and/or GL-7^+^. DZ and LZ cells were gated as CXCR4^hi^ CD86^lo^ and CXCR4^lo^ CD86^hi^. BM transplanted cells in chimeric mice were distinguished by staining cell suspensions with GC markers, as indicated, in addition to the congenic markers CD45.1 and CD45.2 (eBioscience and BioLegend, respectively). Antigen-specific cells in PE- and NP-KLH immunization experiments were detected by staining cell suspensions with GC markers along with PE, PE-NP12 or Pe-NP28. Stained cell suspensions were analyzed using a CytoFlex flow cytometer (Beckman Coulter). For RNA-sequencing, cells were stained for negative markers (dump-: CD4^-^, CD8^-^, GR-1^-^, F4/80^-^) in addition to GC markers and sorted directly into 40μl Lysis/Binding buffer (Life Technologies) using a FACS ARIA II or FACS ARIA III (Becton Dickinson) and immediately frozen on dry ice.

### Transcriptomic analysis

mRNA from 2-5 x 104 sorted cells was captured with 12 μl of Dynabeads oligo(dT) (Life Technologies), washed, and eluted at 85°C with 6.5 ml of 10 mM Tris-HCl (pH 7.5). A bulk adaptation of the MARS-Seq protocol (Jaitin et al., Science 2014, Keren-Shaul et al., Nature Protocols, 2019) was used to generate RNA-Seq libraries for expression profiling of AID-Mettl3^fl/+^ and AID-Mettl3^fl/fl^ GC cells 1 and 2 weeks after immunization or WT and Igf2bp3^-/-^ GC cells 2 weeks after immunization. A minimum of three biological replicates were included in each population. Briefly, mRNA from each sample was barcoded during reverse transcription and pooled. Following Agencourt Ampure XP beads cleanup (Beckman Coulter), the pooled samples underwent second strand synthesis and were linearly amplified by T7 in vitro transcription. The resulting RNA was fragmented and converted into a sequencing-ready library by tagging the samples with Illumina sequences during ligation, RT, and PCR. Libraries were quantified by Qubit and TapeStation as well as by qPCR for ACTB gene as previously described (Jaitin et al., 2014; Keren-Shaul et al., 2019). Sequencing was done on a Nextseq 75 cycles high output kit (Illumina; paired end sequencing).

Alignment and differential expression analysis was performed using the UTAP pipeline (Kohen et al., 2019). Reads were trimmed using Cutadapt and mapped to the Mus_musculus genome (UCSC mm10) using STAR (Dobin et al., 2013) v2.4.2a with default parameters. The pipeline quantifies the genes annotated in RefSeq (extended by 1,000 bases toward the 5’ edge and 100 bases in the 3’ direction). Counting of sequenced reads was done using htseq-count (Anders et al., 2014); union mode). Genes having a minimum of five UMI-corrected reads in at least one sample were included in the analysis. Normalization of the counts and differential expression analysis was performed using DESeq2 (Love et al., 2014) with the following parameters: betaPrior = True, cooksCutoff = FALSE, independentFiltering = FALSE. Raw P values were adjusted for multiple testing using the procedure of Benjamini and Hochberg.

Gene annotation and pathway analysis was carried out using Metascape (Zhou et al., 2019). Gene set enrichment analysis was carried out using GSEA 3.0 with GSEA-preranked tool (Mootha et al., 2003; Subramanian et al., 2005). Gene names were converted to human gene symbols and GSEA run with default parameters (1000 permutations). The Molecular Signature Database hallmark gene sets were used to perform pathway enrichment analysis using a hypergeometric distribution. Gene sets for GC formation and GC B cell selection were generated from GSE110669 and GSE98778 respectively, based on the 200 most significantly upregulated genes in each condition. Dataset used for expression profiling in GC and DZ and LZ cells was GSE93554.

### RT-qPCR

RNA was extracted using Dynabeads™ mRNA DIRECT™ Purification Kit (Invitrogen) and converted to cDNA using either qScript (Quantabio) or Super Script III (Invitrogen) enzyme according to manufaturer’s instructions with a mix of random and poly A-specific primers. For qPCR reactions a QuantStudio 5 Real-Time PCR system and Fast SYBR Green Master Mix (Applied Biosystems) were used. Primers are listed in Table S2.

### In vitro proliferation assays

Single cell suspensions were obtained from mouse spleens and stained with CellTrace-Violet (Molecular Probes) according to manufacturer’s instructions. Cells were seeded in a 96 well plate at 1.5 x 10^6^/ml and stimulated with LPS, αIgM or αCD40 at concentrations ranging from 10ng to 10μg per ml for three days. Cell proliferation was analyzed by flow cytometry.

### ELISA for total immunoglobulin titers

Serum was collected from mice and titers of IgM, IgG1 and IgG2 antibodies were determined by direct ELISA. Diluted serum was used to coat 96 well plates by incubating overnight at 4oC. Relative quantities of antibodies were detected using HRP-conjugated anti-mouse IgM, IgG1 and IgG2 (Abcam) conjugated to horseradish peroxidase.

### Immunohistochemistry

Fixed single popliteal LNs were frozen in OCT and cut into 10 μm sections. Slides were blocked and incubated with fluorescently labeled antibodies. Anti-IgD and GL-7 antibodies were from Biolegends. Germinal centers were identified as IgD^-^ GL-7^+^ areas. Images were captured using a 20X objective. For imaging specifications see smFISH section.

### Single-cell Igh sequencing

Single popliteal LNs were harvested from AID-Mettl3^fl/fl^ mice and processed for flow cytometric analysis. Cell suspensions were stained for negative markers (dump-: CD4^-^, CD8^-^, GR-1^-^, F4/80^-^) and gated as dump^-^ B220^+^ GL-7^+^ CD38^-^ FAS^+^ IgG1^+^ or dump^-^ B220^+^ GL-7^+^ CD38^-^ FAS^+^ IgG1^+^ IgL^+^ for isolation of VH186.2-expressing cells. Cell sorting was performed using a FACS Aria II cell sorter (BD Bioscience). For total V(D)J sequencing of Ighγ1 heavy chains, GC-derived B cells were sorted into 96-well plates containing lysis buffer (PBS containing 3 U μl–1 ribonuclease inhibitor and 10 mM dithiothreitol). For sequencing of Ighγ1 heavy-chains from VH186.2-expressing cells, GC-derived B cells were sorted in bulk into lysis/binding buffer (Life technologies)

### Single cell total IgG sequencing and clonal analysis

Complementary DNA from single cells was generated using random primers (NEB) as previously described. Ighγ1 heavy-chain sequences were amplified in a nested PCR using primers for the Ighγ1 constant region together with a mix of primers for the variable regions (von Boehmer et al., 2016). Complementary DNA from bulk sorting was produced using qScript cDNA Synthesis Kit (QuantaBio). VH186.2 (V1-72*01) Ighγ1 heavy-chain sequences were amplified in a nested PCR using primers for the Ighγ1 constant region together with a specific primer for VH186.2 (Mayer et al., 2017)(Table S1) and using high-fidelity Q5 polymerase (NEB). Amplified products were cloned (CloneJET PCR Cloning Kit, ThermoScientific), sequenced and analyzed.

PCR products were sequenced by sanger sequencing and CDR3 regions were analyzed by aligning Ig Fasta sequences against the IMGT mouse heavy gene database (September 2017) using IgBlast (v.1.7.0) (Ye et al., 2013). Sequence alignment was performed using SnapGene software (GSL Biotech). For a schematic representation of total Ighγ1-sequence results, sequences derived from single cells were clustered according to their CDR3 region (Identical V, D and J segment represents one clone) and presented as part of total sequenced cells. Primer-derived mutations were excluded from the analysis. For schematic representation of VH186.2-sequence results, sequences were clustered according to presence/absence of W33L and K59R mutations and presented as part of total sequenced cells.

### Isolation of B cell mRNA

Resting B cells were isolated from mouse spleens by forcing tissue through 70 μm mesh into PBS. B cells were purified from cell suspensions using αCD43 magnetic beads (MACS) according to manufacturer’s instructions (Militenyi Biotec). B cell purity of minimum 96% was confirmed by staining for B220 and analyzed by flow cytometry. Isolated B cells were stimulated with LPS (5mg/ml) for one day in vitro at 1.5 x 106 cells per ml in RPMI complete medium (10% FCS, Pen/Strep, L-glutamine, HEPES, β-mercaptoethanol). Total RNA was isolated using Trizol reagent (invitrogen) and polyadenylated RNA was purified twice using oligo(dT) beads (Life Technologies).

### M^6^A-Seq and Analysis of m6A Methylation Sites

M^6^A immunoprecipitation and RNA-seq library preparation was performed as previously described (Garcia-Campos et al., 2019). Three biological replicates of control and METTL3-deleted samples were included. Reads were aligned to mouse mm9 genome using STAR aligner (Dobin et al., 2013). A previously published approach was used to identify putative m6A sites (Schwartz et al., 2014). Each site was assigned a peak over input (POI) and peak over median (POM) scores representing the fold change of enrichment in the peak region over the corresponding region in the input fraction or over the median coverage in the gene, respectively, as in Schwartz et al (Schwartz et al., 2014). For identification of sites present in the control sample but absent in the METTL3-deleted sample, we performed two separate t-tests to assess whether the POM and/or the POI scores differed significantly (P < 0.05) between control and METTL3-deleted samples. A peak was determined to be METTL3-specific if the mean POI and mean POM scores (across the triplicates) in the WT samples were higher than their counterparts in the METTL3-deleted samples and if at least one of the two associated P values was significant. For estimating the global methylation levels per sample (Fig. S7C-D) we calculated POI and POM scores for a pre-assembled set of 16,321 consistently identified sets of m6A sites and a set of 30,115 control sites (Garcia-Campos et al., 2019).

### smFISH - hybridizations and imaging

LNs from AID-Mettl3^fl/+^ and Mettl3^fl/fl^ mice were extracted and fixated in 4% paraformaldehyde in ultra clean water (Sigma-Aldrich). 5μm (exon only slides) or 8μm (exon and intron slides) slices were cryo-sectioned, embedded on Poly-L-lysine treated coverslips and washed with ice cold 70% ethanol. Coverslips were then placed in a 6-well plate and mounted with hybridization mix overnight. Hybridization mix consisted of 10% Dextran sulfate, 15% formamide, E. coli tRNA (1mg/ml), SSC 2X, 0.02% BSA, 2mM Vanadyl Ribonucleoside complex with both probes (1:3000) according to previously described protocol (Itzkovitz et al., 2011).

Probe libraries were designed and constructed as described in Raj et al (Raj et al., 2008). Myc intron and exon libraries consisted of 48 probes of 20bp length, complementary to the intronic sequence or coding sequence, respectively. Ki67 libraries consisted of 96 probes (Table S3). Hybridizations were done overnight with two to three differentially labelled probes using Cy5, Alexa594 and TMR fluorophores. A488-conjugated antibody for B220 (BioLegend) was added to the hybridization mix and used for protein immunofluorescence. 4,6-diamidino-2-phenylindole dye for nuclear staining was added during the washes. Images were taken with a Nikon Ti-2 inverted fluorescence microscope equipped with a ×100 oil-immersion objective and a Photometrics Prime BSI sCMOS camera using NIS-Elements AR software (v5.20.01)(Nikon). The image-plane pixel dimension was 0.13 μm. Quantification was done on stacks of optical sections, in which not more than a single cell was observed. Dots were automatically detected using Imaris software (v. 9.5.1, Oxford Instruments). Automatic threshold selection was manually verified and corrected for errors. Background dots were detected according to size and by automatically identifying dots that appear in more than one channel (typically <1% of dots) and were removed. Such dots occasionally appeared in the surrounding non-B cells but were rare in the B220^+^ B cells. Bleed-through of transcript signal between channels was minimal. Cell segmentation was carried out manually on a maximal projection of the A488 channel.

### Statistical analysis and reproducibility

Statistical significance was determined using Graphpad Prism v.7.0 using the tests indicated in each figure.

