## Supplementary Material for "B Cell Division Capacity in Germinal Centers Depends on Myc Transcript Stabilization Through m^6^A mRNA Methylation and IGF2BP3 Functions"

**Figure S1**

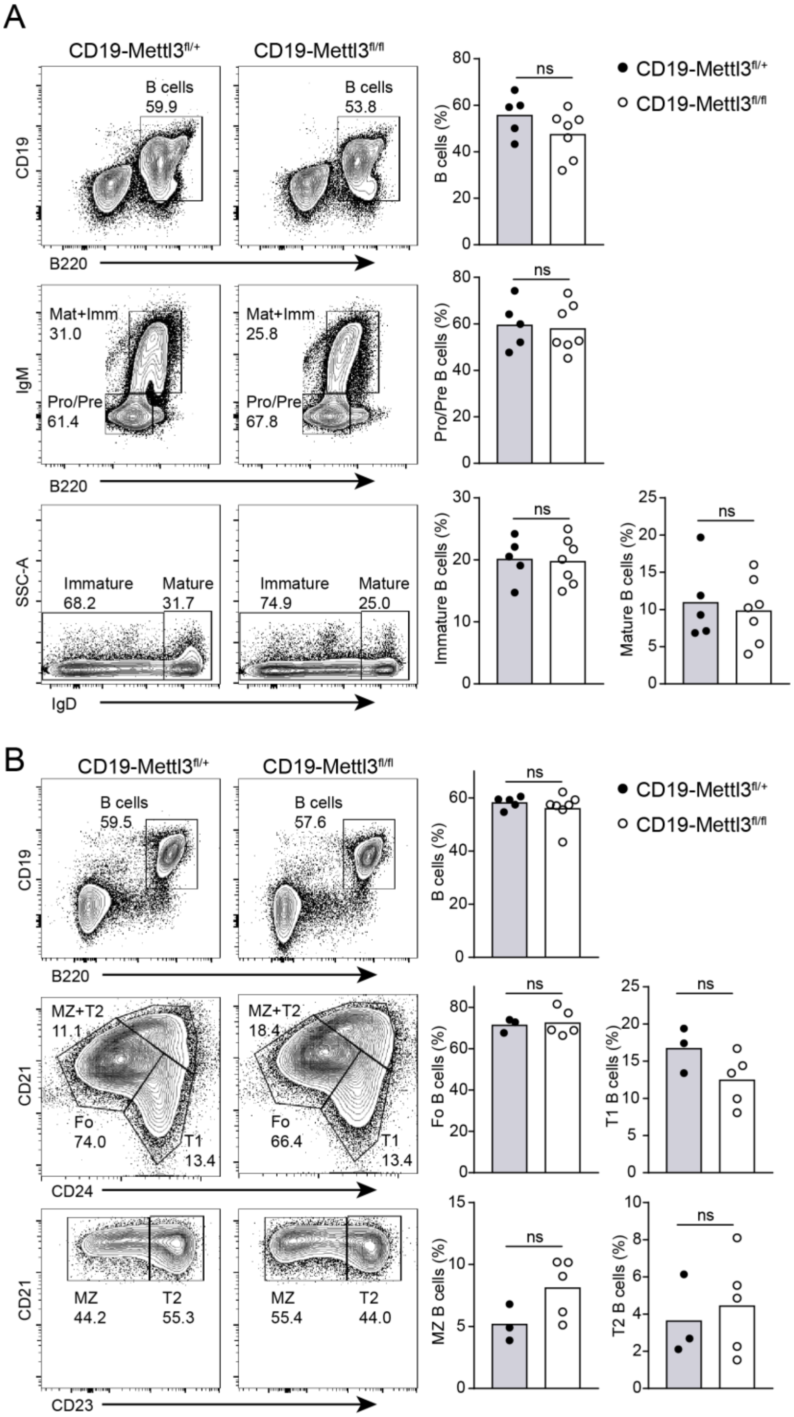


**Figure S1. B cell development in control and CD19-Mettl3$^{fl/fl}$ mice.** (**A, B**) Flow cytometer analysis of B cell developmental stages in the BM (A) and spleen (B). The size of each population is summarized in the right panels. Pooled data from two to three experiments each with one to three mice per condition. Two-tailed student's *t* test. NS, not significant.

### Figure S2

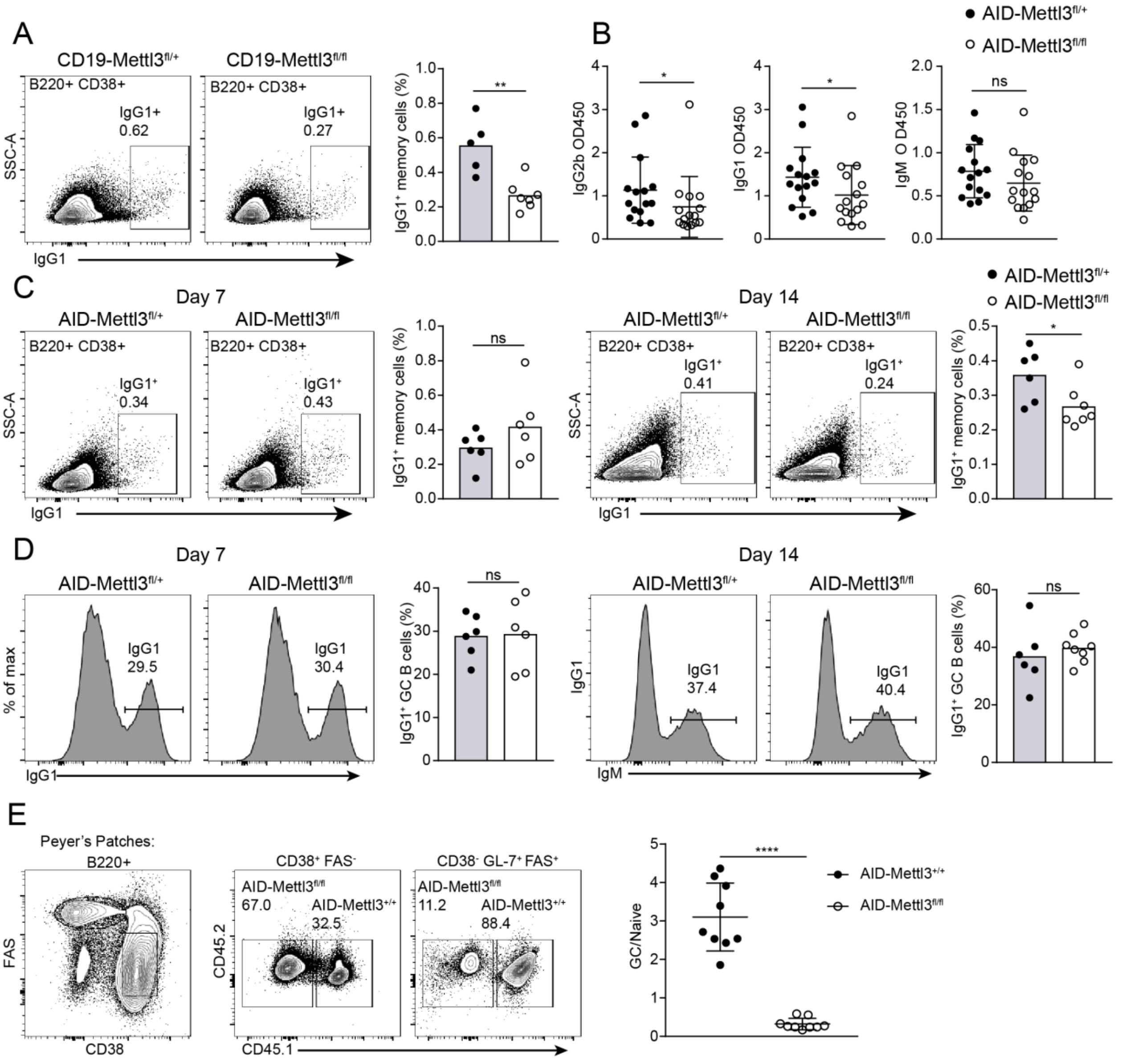
A
CD19-Mettl3fl/+
CD19-Mettl3fl/fl
B220+ CD38+
IgG1+
0.62
0.27
SSC-A
IgG1
IgG1+ memory cells (%)
**
B
AID-Mettl3fl/+
AID-Mettl3fl/fl
IgG2b OD450
IgG1 OD450
IgM O D450
*
ns
C
Day 7
Day 14
AID-Mettl3fl/+
AID-Mettl3fl/fl
0.34
0.43
0.41
0.24
D
% of max
IgG1
29.5
30.4
37.4
40.4
IgG1+ GC B cells (%)
IgM
E
Peyer's Patches:
B220+
FAS
CD38
CD38+ FAS-
CD38- GL-7+ FAS+
AID-Mettl3+/+
67.0
32.5
11.2
88.4
CD45.2
CD45.1
GC/Naive
****

**Figure S2. METTL3 is required for memory B cell formation but not for isotype class switch recombination.** (**A**) CD19-Mettl3$^{fl/+}$ or CD19-Mettl3$^{fl/fl}$ mice were immunized in the footpad with NP-KLH. After 7 days, the relative numbers of CD38$^{+}$ IgG1$^{+}$memory B cells were examined by flow cytometry. Data pooled from three experiments with 5-7 mice per condition in total. (**B**) Serum immunoglobulin titers in AID-Mettl3$^{fl/+}$ or AID-Mettl3$^{fl/fl}$ mice. 15 mice per condition, student's paired *t* test between age-matched mice was used to determine significance. (**C**) AID-Mettl3$^{fl/+}$ or AID-Mettl3$^{fl/fl}$ mice were immunized with NP-KLH in the footpad. After 7 and 14 days, the relative numbers of CD38$^{+}$ IgG1$^{+}$ memory B cells were examined by flow cytometry. Pooled data from three experiments each with 6-7 mice per condition in total. (**D**) The frequency of IgG1$^{+}$ B cells in GCs of AID-Mettl3$^{fl/+}$ or AID-Mettl3$^{fl/fl}$ 7 and 14 days after immunization. (**E**) Chimeric mice were generated by transfer of a mix of cells containing 70 % control BM cells (AID$^{cre/+}$) and 30% AID-Mettl3$^{fl/fl}$ BM cells. The frequency of GC cells in Peyer's patches of control and AID-Mettl3$^{fl/fl}$ control were examined by flow cytometry. Pooled data from two to three experiments with 2-5 mice per condition. Column heights represent mean (A-D) or bars represent mean+SD (E). Unpaired student's *t* test was used to determine significance. *$P < 0.05$; **$P < 0.01$; ***$P < 0.005$; ****$P < 0.001$. NS, not significant.

**Figure S3**

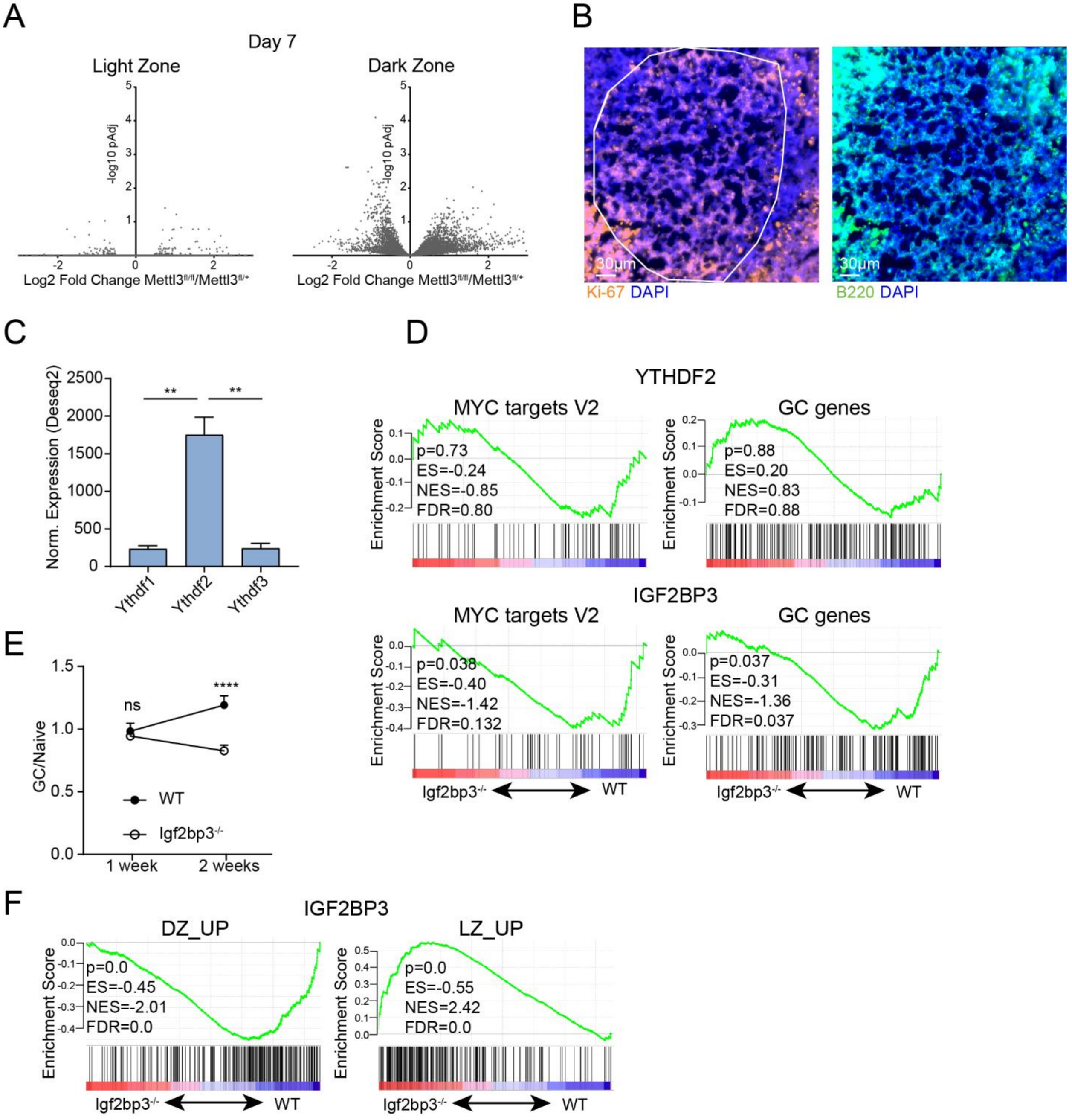
A
Day 7
Light Zone
Dark Zone
-log10 pAdj
Log2 Fold Change Mettl3fl/fl/Mettl3fl/+
B
30µm
Ki-67 DAPI
B220 DAPI
C
Norm. Expression (Deseq2)
Ythdf1
Ythdf2
Ythdf3
D
YTHDF2
MYC targets V2
p=0.73
ES=-0.24
NES=-0.85
FDR=0.80
GC genes
p=0.88
ES=0.20
NES=0.83
FDR=0.88
IGF2BP3
MYC targets V2
p=0.038
ES=-0.40
NES=-1.42
FDR=0.132
GC genes
p=0.037
ES=-0.31
NES=-1.36
FDR=0.037
Enrichment Score
Igf2bp3-/-
WT
E
GC/Naive
ns
****
WT
Igf2bp3-/-
1 week
2 weeks
F
IGF2BP3
DZ_UP
p=0.0
ES=-0.45
NES=-2.01
FDR=0.0
LZ_UP
p=0.0
ES=-0.55
NES=2.42
FDR=0.0
Enrichment Score
Igf2bp3-/-
WT

**Figure S3. METTL3 regulates the DZ compartment in GC B cells.** (**A**) Additional supporting results from the experiments in Fig. 3B and corresponding text. Differential gene expression of control and Mettl3-deficient cells 7 days after immunization with NP-KLH. (**B**) Analysis of in GC B cells by smFISH. GCs were identified by staining with B220 antibody and Ki-67 smFISH probe. (**C**) Analysis of YTHDF1/2/3 expression in RNAseq data of GC B cells based on dataset GSE98778. (**D**) Additional supporting results from the experiments shown in Fig. 6C. GSEA of GC B cells derived from Ythdf2- and Igf2bp3-deficient mice and control mice. (**E**) Chimeric mice generated by BM transfer of 30% WT and 70% Igf2bp3$^{-/-}$ cells were analyzed for the fraction of each cell type in the naïve and GC B cell compartments 1 and 2 weeks after immunization with NP-KLH. Pooled data from 3 experiments, each with three to four mice. Each datapoint represents mean+SD. (**F**) Additional supporting results from the experiment shown in Fig. 6C and corresponding text. GSEA plots of DZ and LZ gene signatures in Igf2bp3$^{-/-}$ vs. WT GC B cells. Statistical significance in C and E was determined One-way ANOVA with Sidak's multiple comparisons test. ***$P$ < 0.005, NS, not significant.

**Figure S4**

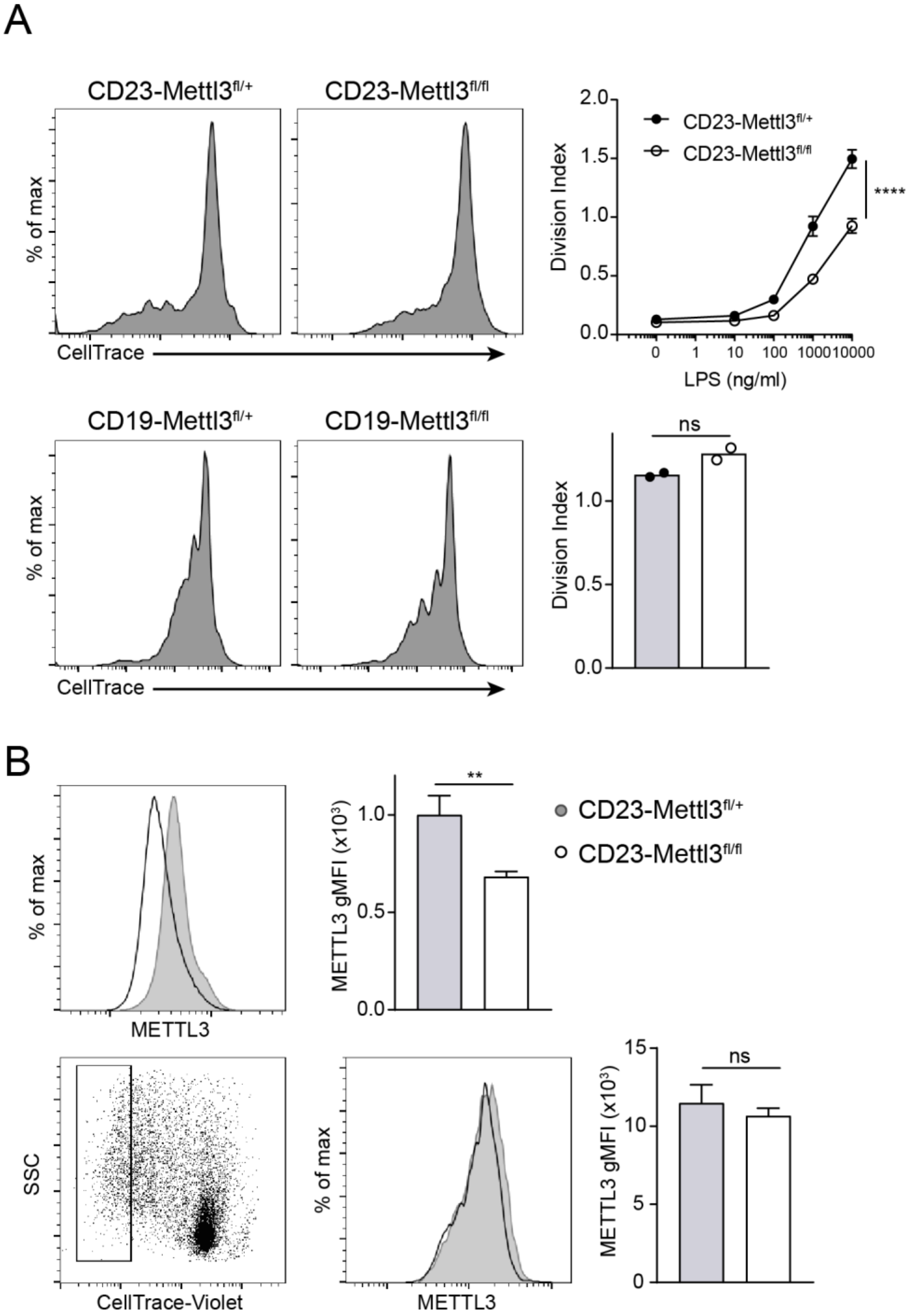
A
CD23-Mettl3fl/+
CD23-Mettl3fl/fl
% of max
CellTrace
Division Index
LPS (ng/ml)
0
1
10
100
1000
10000
****
CD19-Mettl3fl/+
CD19-Mettl3fl/fl
ns
B
METTL3
METTL3 gMFI (x10³)
**
SSC
CellTrace-Violet

**Figure S4. In vitro LPS-induced proliferation and METTL3 expression of control and Mettl3-deficient B cells.** (**A**) Isolated splenic cells from CD19 or CD23-Mettl3$^{fl/+}$ and control mice were stained with CellTrace-Violet and stimulated *in vitro* with LPS at increasing concentrations. After 3 days cells were analyzed by flow cytometry. Representative plots from 2-3 experiments, each with two to three replicates. Mean+SD (top) or mean (bottom) are shown. (**B**) Fresh splenic B cells (top) or actively dividing B cells (bottom) from mice in a were stained intracellularly for expression of METTL3. Two-way ANOVA (A, top) and two-tailed Student's t-test (A, bottom and B). Pooled data from 2 experiments each with three replicates. Mean+SD is shown. **= $P < 0.01$; ****= $P < 0.001$. NS, not significant.

**Figure S5**

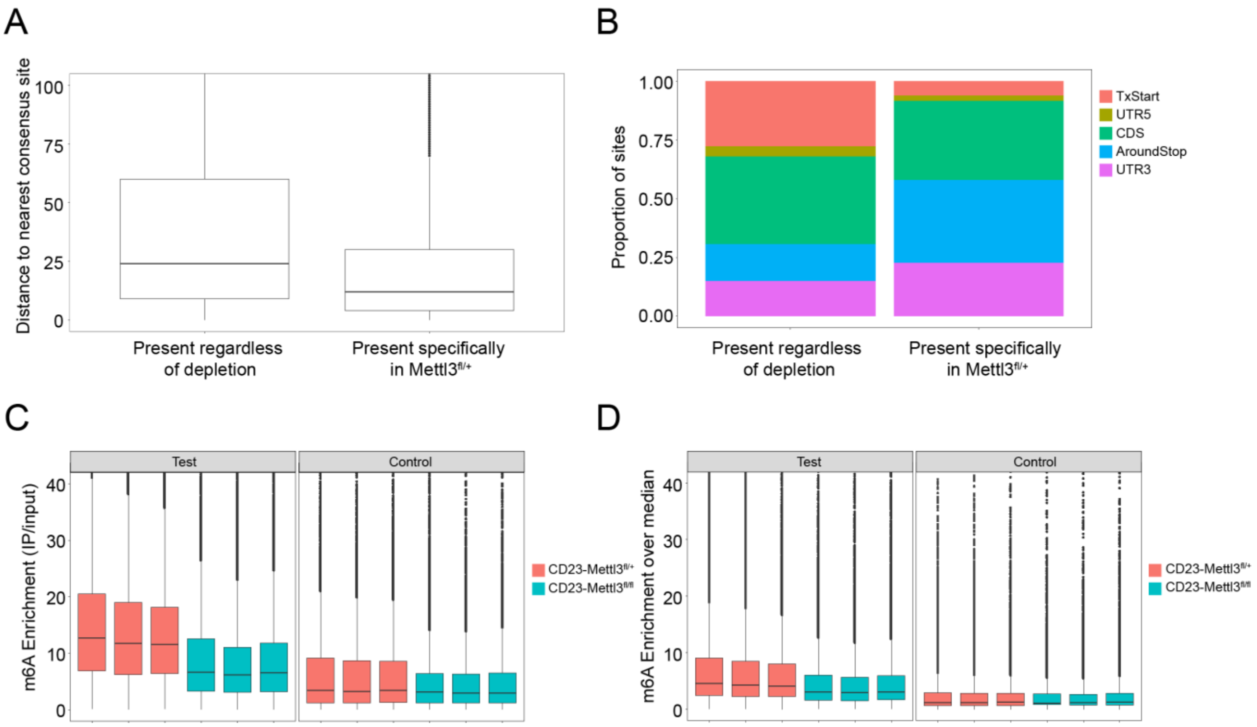


**Figure S5. B cells derived from Mettl3-deficient mice have decreased levels of m$^6$A-methylation.** (**A**) Analysis of m$^6$A abundance in B cells that were stimulated with LPS *in vitro*. Distributions of distance from the nearest consensus site of sites that are present only in control samples or both control and Mettl3-deficient B cells. (**B**) Distribution of the sites in (A) grouped across five gene segments as defined in Dominissini et al., 2012. (**C, D**) Calculated Peak over Input (C) and Peak over Median (D) scores for control and Mettl3-deficient B cells.

**Table S1**

| Antigen | Fluorophore | Clone | Manufacturer | Concentration, µg/ml |
|---|---|---|---|---|
| B220 | APC-Alexa Flour 780 | RA3-6B2 | eBioscience | 1 |
| B220 | V500 | RA3-6B2 | BD | 1 |
| B220 | e450 | RA3-6B2 | eBioscience | 1 |
| B220 | APC | RA3-6B2 | BioLegend | |
| CD138 | BV605 | 281-2 | Biolegend | 0.5 |
| CD38 | Alexa Flour 700 | 90 | eBioscience | 1 |
| F4/80 | APC-Alexa Flour 780 | BM8 | eBioscience | 1 |
| FAS | PE-Cy7 | Jo2 | BD | 0.33 |
| FAS | Alexa Fluor 647 | Jo2 | BD | 0.33 |
| GL-7 | Alexa Flour 647 | GL7 | Biolegend | 2.5 |
| GL-7 | FITC | FR70 | Biolegend | 2.5 |
| GL-7 | PerCP-Cy5.5 | GL7 | Biolegend | 1 |
| Gr-1 | APC-Alexa Flour 780 | RB6-8C5 | eBioscience | 1 |
| Igλ | APC | RML-42 | Biolegend | 1 |
| IgA | PE | mA-6E1 | eBioscience | 1 |
| CD24 | PE-Cy7 | M1/69 | BioLegend | 0.5 |
| CD21 | FITC | 7E9 | BioLegend | 0.5 |
| CD23 | PE | B3B4 | BioLegend | 0.5 |
| IgG1 | FITC | RMG1-1 | BioLegend | 0.5 |
| IgG1 | BV510 | RMG1-1 | BioLegend | 0.5 |
| CD86 | PE | GL-1 | BioLegend | 0.13 |
| CD86 | Alexa Fluor 647 | GL-1 | BioLegend | 0.4 |
| CD86 | FITC | GL-1 | BioLegend | 0.4 |
| CXCR4 | BV421 | L276F12 | BioLegend | 0.5 |
| IgD | Alexa Fluor 647 | 11-26C.2A | BioLegend | 0.5 |
| IgM | PerCP-eFluor 710 | 11/41 | Invitrogen | 0.5 |
| CD19 | Pacific Blue | 6D5 | BioLegend | 0.5 |
| Rabbit METTL3 | - | PolyAb | ProteinTech | 0.5 |
| Goat Anti-Rabbit | Alexa Fluor 488 | polyAb | Invitrogen | 0.13 |

**Table S1: Antibodies used in flow cytometry**

**Table S2**

| List of primers for RT-qPCR | |
|---|---|
| **Primer Name** | **Primer sequence (5'-3')** |
| Igγ1 constant region, Outer | GGAAGGTGTGCACACCGCTGGAC |
| Igγ1 constant region, Inner | GCTCAGGGAAATAGCCCTTGAC |
| VH186.2 | CTAGTAGCAACTGCAACCGGTGTACATTCTCAGGTGCAGCTGCAGGAGTC |
| Hprt fwd | TATGGCGACCCGCAGCCCT |
| Hprt rev | CATCTCGAGCAAGACGTTCAG |
| Ubc fwd | GCCCAGTGTTACCACCAAGA |
| Ubc rev | CCCATCACACCCAAGAACA |
| Gapdh fwd | TTGATGGCAACAATCTCCAC |
| Gapdh rev | CGTCCCGTAGACAAAATGGT |
| Mettl3 fwd | GAGTTGATTGAGGTAAAGCGAGG |
| Mettl3 rev | GGAGTGGTCAGCGTAAGTTACA |
| Myc fwd | AAACGACAAGAGGCGGACACAC |
| Myc rev | AAAGCTGCGCTTCAGCTCGTTC |
| Myc_m6A_fwd | CTGTCCATTCAAGCAGACGA |
| Myc_m6A_rev | TCCAGCTCCTCCTCGAGTTA |

**Table S2: Primers for RT-qPCR**
